# A full-length isoform atlas of the non-failing and failing human heart

**DOI:** 10.64898/2026.11.13.766598

**Authors:** Bo Ye, Mehwish Noureen, Funan He, Lakshman Lingamgunta, Siyuan Zheng, Faqian Li, Xiaojing Wang

**Author notes:** Correspondence should be addressed to: F.L. and X.W and. These authors contributed equally (Bo Ye, Mehwish Noureen). The authors declare no potential conflicts of interest.

## Abstract

Dilated cardiomyopathy (DCM) and ischemic heart disease (IHD) are leading causes of heart failure, yet isoform-level changes in the failing heart remain poorly defined because reference annotations are derived largely from healthy tissues and short-read sequencing cannot reliably reconstruct full-length transcripts. We sequenced six human left ventricles (non-failing, DCM, and IHD; n = 2 each) with PacBio HiFi long reads and curated 48,232 isoforms across 11,760 genes, including 15,535 novel NIC and NNC isoforms. Independent evidence supported most novel structures: 81.7% of isoforms with novel splice junctions had short-read support for all novel junctions, and a cardiac CAGE atlas supported the 5′ ends of 93% of NIC and 86% of NNC isoforms. Novel isoform diversification was concentrated in sarcomere genes and persisted after accounting for gene complexity, expression, sequencing depth, and transcript length. We then augmented GENCODE v43 with 18,637 non-FSM long-read isoforms and re-quantified two independent short-read cohorts (MAGNet and GSE116250). For most genes carrying a transcript that changed more than 2-fold in DCM, the gene itself changed less than 2-fold, so these changes are not apparent at the gene level. Of 626 transcripts differentially expressed in both cohorts, 98% changed in the same direction, including all 100 long-read isoforms among them. The resource is available through CardioIsoMap, an interactive isoform-centric portal (https://cardioisomap.streamlit.app/). This study provides a comprehensive isoform-resolved reference of the human heart in health and disease.

## Introduction

Heart failure (HF) remains a leading cause of morbidity and mortality worldwide^1^. Two of its principal etiologies are dilated cardiomyopathy (DCM) and ischemic heart disease (IHD), which have distinct but partially overlapping mechanisms. DCM is dominated by genetic and structural remodeling of the sarcomere and cytoskeleton, frequently progressing to chamber enlargement and reduced ejection fraction, whereas IHD follows coronary ischemic injury and the adverse remodeling, fibrosis, and inflammatory signaling that accompany cardiomyocyte loss^2-4^.

Alternative splicing and alternative transcript start and end usage are pervasive in the heart and are central to cardiomyocyte structure, contractility, and metabolism^5-7^. Splicing dysregulation is increasingly recognized as a driver of cardiac dysfunction. Mutations in RNA-binding proteins such as RBM20 cause DCM through aberrant splicing of sarcomere transcripts^8^, and broad reductions in splicing efficiency affecting sarcomere genes including TNNT2, TNNI3, MYH7, and FLNC have been reported in IHD myocardium^9^. These observations argue that isoform-level changes carry disease information that gene-level summaries cannot capture.

Despite extensive transcriptomic profiling of failing myocardium, most studies have operated at the gene level and on reference annotations derived predominantly from healthy tissue^10-12^. This limitation is partly methodological. Short-read RNA sequencing infers rather than observes full-length transcripts, and its read length together with reliance on junction-spanning reads prevents unambiguous reconstruction of novel exon combinations and accurate resolution of transcript boundaries^13,14^. Benchmarking studies confirm that short-read methods recover only a fraction of events such as intron retention, with poor concordance across tools^15^. Long-read sequencing instead captures full-length molecules in single reads and is therefore uniquely suited to building isoform-resolved annotations. Long-read transcriptomic atlases have been generated for cell lines (LRGASP)^16^, the human brain^17^, and a panel of GTEx tissues that included postmortem atrial and left-ventricular samples^18,19^. In the heart, long reads have resolved isoforms disrupted by RBM20 mutations in cardiomyocytes^20^, cell-type-specific isoform usage in single nuclei from normal and failing left ventricles^21^, and isoform shifts in sarcomere genes in left ventricles from patients with dilated and ischemic cardiomyopathy^22^. These studies show that the heart expresses many unannotated isoforms, but two questions remain open: how much of the novel content in a cardiac long-read catalog is supported by independent evidence, and whether isoform changes detected in small long-read cohorts reproduce in large patient cohorts.

Here, we characterize full-length transcript isoforms in human left ventricles from non-failing, DCM, and IHD hearts using PacBio HiFi Iso-Seq. We curate the resulting catalog and independently evaluate novel splice junctions, exons, and transcript ends using short-read RNA-seq and cardiac CAGE data. We then incorporate the curated isoforms into GENCODE and re-quantify two independent short-read cohorts comprising more than 400 left ventricles (MAGNet and GSE116250) to assess population-level expression and replication of disease-associated isoform changes. Our results reveal widespread disease-associated isoform alterations, many involving previously unannotated transcripts, with directional concordance between DCM and ischemic cardiomyopathy and frequent absence of corresponding gene-level changes. We make this resource publicly available through CardioIsoMap.

## Results

### A full-length long-read isoform atlas of the human left ventricle

We performed barcoded PacBio HiFi long-read RNA sequencing on six human left-ventricular tissue samples representing non-failing hearts (NFH, n = 2), dilated cardiomyopathy (DCM, n = 2), and ischemic heart disease (IHD, n = 2) (**Fig. 1a; Supplementary Table 1**). Raw subreads were processed through the Iso-Seq workflow into full-length non-chimeric (FLNC) reads, aligned to GRCh38, and collapsed into non-redundant isoforms (**Methods, Supplementary Figure 1**). The dataset comprised approximately 2.4 million FLNC reads (**Supplementary Table 1**). Classifying the collapsed isoforms against GENCODE v43^23^ with SQANTI3^24,25^ yielded an initial catalog of 62,673 isoforms, comprising 29,595 full-splice-match (FSM), 6,076 incomplete-splice-match (ISM), 12,045 novel-in-catalog (NIC), and 13,656 novel-not-in-catalog (NNC), along with 329 fusion, 398 intergenic, 288 antisense, and 286 genic.

**Figure 1.**
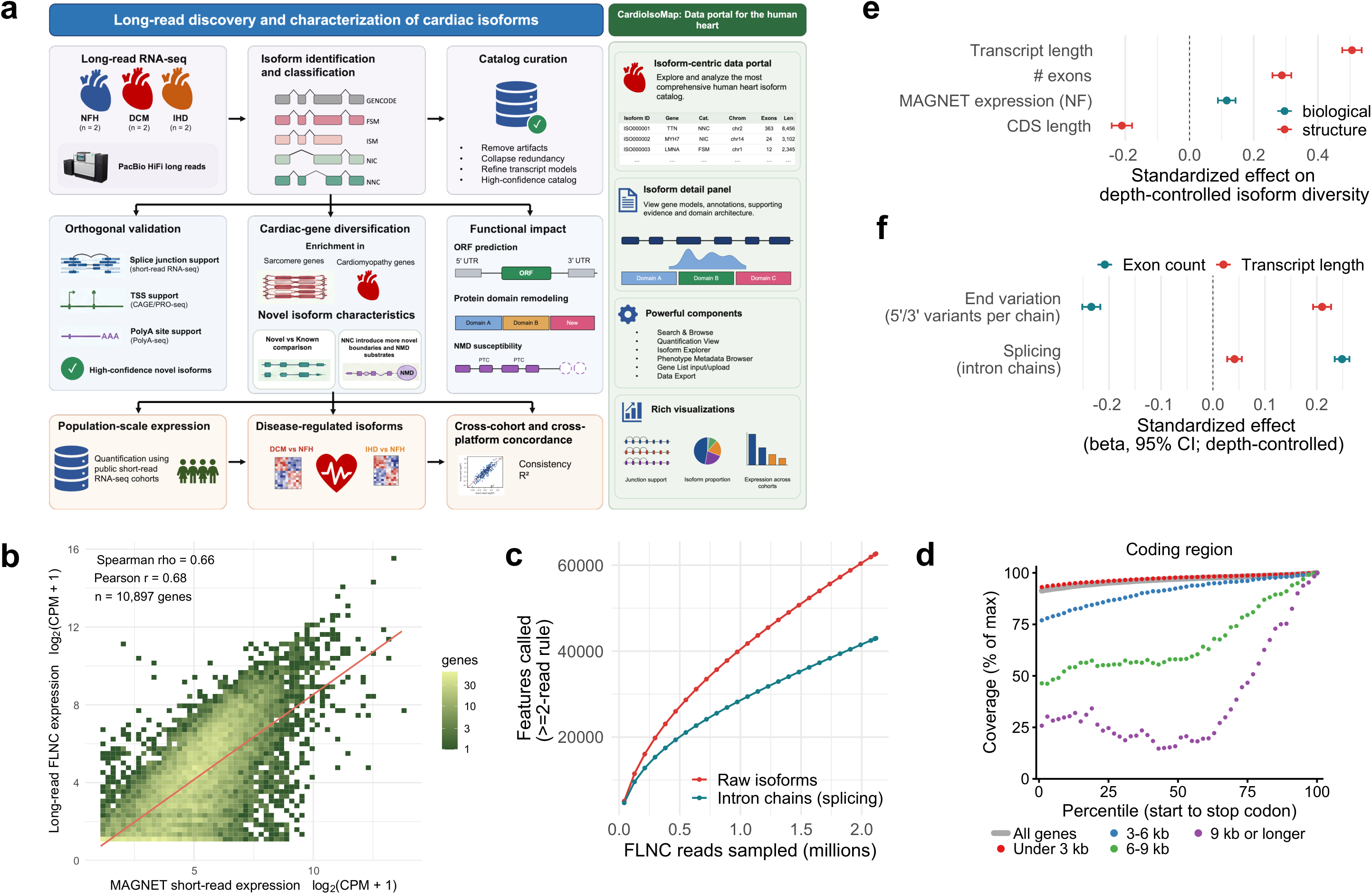
A long-read isoform atlas of the human heart. **a.** Study overview. PacBio HiFi long-read RNA sequencing of non-failing (NFH), dilated cardiomyopathy (DCM), and ischemic heart disease (IHD) hearts (n = 2 per group). Full-length transcripts were identified, classified, and curated to generate a high-confidence cardiac isoform catalog and the CardioIsoMap data portal. **b.** Correlation between long-read FLNC-derived and MAGNet short-read-derived gene expression estimates (log2(CPM + 1); n = 10,897 genes). **c.** Rarefaction curves showing the accumulation of isoforms and unique intron chains with increasing FLNC read depth. **d.** Median gene-body coverage of FLNC reads across the coding region (start to stop codon) of each protein-coding gene’s GENCODE v43 Ensembl canonical transcript. Each gene’s coverage was normalized to its own maximum before calculating the median across genes. Only genes with at least 20 reads at their maximum were included. Genes were grouped by coding length: under 3 kb (shorter than the median read), 3-6 kb, 6-9 kb (shorter than 99% of reads), and 9 kb or longer. All genes, n = 8,319; under 3 kb, n = 6,948; 3-6 kb, n = 1,120; 6-9 kb, n = 181; 9 kb or longer, n = 70. **e**. Standardized effects of transcript length, exon number, CDS length and MAGNet expression in non-failing hearts on isoform diversity after rarefaction to 10 FLNC reads per gene (linear model; n = 8,017 genes). Transcript length and exon number were positively associated with isoform diversity. **f.** Structural determinants of transcript diversity (n = 11,303 genes). Exon number was the primary determinant of splice-junction diversity, whereas transcript length predominantly contributed to variation in transcription start and termination sites.

The full-length FLNC depth per isoform demonstrated a broad dynamic range (median 4, mean 33.8, maximum 79,612), which was reproducible across samples (median detection in 3 of 6 samples). Given the dynamic depth, we sought to quantify abundant isoforms while confirming sparse isoforms with orthogonal, cross-platform evidence. As an independent check on our quantification, gene-level expression from long-read data was moderately concordant with independent MAGNet short-read RNA-seq (Spearman ρ = 0.66, n = 10,897 genes; **Fig. 1b**), indicating agreement between platforms in relative gene abundance and supporting a workflow in which long reads resolve isoform structure while short reads provide robust quantification.

### Detection depth and gene structure shape long-read isoform diversity

Rarefaction of FLNC reads showed that gene discovery saturated rapidly (**Supplementary Figure 2**), whereas isoform discovery continued to climb without reaching a plateau: across the full subsampling range, the catalog was still gaining roughly 1,800 new isoforms and 1,200 new intron chains per additional 100,000 reads (**Fig. 1c**).

To assess the completeness of long-read coverage, we examined FLNC read coverage at 100 percentile positions along the GENCODE v43 canonical transcript of each protein-coding gene, over its coding region and over the full transcript (**Fig. 1d, Supplementary Figure 3**, **Supplementary Table 2**). Coverage was close to uniform across the coding regions of most genes (87% of its maximum at the start codon across all 19,978 coding regions, **Supplementary Figure 3b**; median gene, 91%) but declined once coding regions exceeded the read length (median 3.2 kb; 99% of reads shorter than 9.1 kb): in the median gene, the start codon reached 93% of the maximum for coding regions under 3 kb, 77% for 3-6 kb, 46% for 6-9 kb and 26% for 9 kb or longer (**Fig. 1d**). Along the full transcript, coverage also declined toward both ends (median gene, 48% at the first and 11% at the last percentile, **Supplementary Table 2**; all genes summed, 62% and 42%, **Supplementary Figure 3a**), consistent with incomplete capture of transcript termini and with reads ending at upstream poly(A) sites, a pattern also reported for ONT direct RNA sequencing^26,27^. Profiles were similar in all six hearts (**Supplementary Figure 3c**). Molecules much longer than the typical read were therefore infrequently captured in full, under-representing the 5′ portion of long coding regions and potentially contributing to ISM artifacts, which we test directly below.

Per-gene isoform count is confounded by sequencing depth. In a multivariable negative-binomial model of 10,840 genes, FLNC read depth was the dominant predictor of isoform count (ρ = 0.76; standardized β = 0.68), and the univariate association with MAGNet expression (ρ = 0.48) disappeared after accounting for depth (partial ρ = -0.05; **Supplementary Figure 4**), indicating that it reflects increased transcript sampling among highly expressed genes. To remove the depth effect directly, we rarefied each gene with ≥10 FLNC reads to a common depth of 10 reads (n = 8,017 genes), which made isoform diversity much less depth-dependent (ρ = 0.23, down from 0.61). After rarefaction, transcript length and exon count showed the strongest positive associations with isoform counts (standardized β = 0.51 and 0.29), CDS length was negatively associated (β = -0.21), and independent MAGNet expression showed only a weak association (β = 0.12, **Fig. 1e**).

Collapsing isoforms with identical intron chains reduced the 48,232 curated isoforms by 32%, leaving 32,703 distinct intron chains. This indicates that a substantial proportion of isoform diversity reflects alternative 5′ and 3′ ends rather than differences in splicing patterns^6^. In depth-controlled models of the two components (n = 11,303 genes), exon count was the dominant structural predictor of splicing (β = 0.25, 95% CI [0.24, 0.26]) but was negatively associated with end variation (β = -0.23, 95% CI [-0.25, -0.22]), whereas transcript length was strongly associated with end variation (β = 0.21, 95% CI [0.19, 0.23]) but showed little association with splicing (β = 0.04, 95% CI [0.03, 0.06]) (**Fig. 1f**). Thus, after controlling for depth, exon count reflects the combinatorial complexity of splicing^28^, whereas transcript length contributes to isoform diversity through alternative 5′/3′ ends.

### Curation of a high-quality transcriptome reference

The raw isoform catalog contained both *bona fide* transcripts and potential artifacts arising from reverse-transcription template switching, intra-priming, incomplete transcript ends, and splice-site alignment noise. We therefore applied category-aware curation combining SQANTI3 machine-learning classification^24^ with transcript-end support and reproducibility criteria.

FSM isoforms (n = 29,595), which recapitulated the exon-intron structures of annotated transcripts, were retained without additional filtering and accounted for 61.4% of the final reference. Among 6,076 ISMs, 2,495 (41.1%) were retained based on appropriate transcript-end support, including 1,839 5′-covering transcripts, 272 intron-retention isoforms, 362 3′-covering transcripts, and 22 internally supported fragments. The remaining 3,581 ISMs were excluded (**Supplementary Table 3**).

Among NIC, NNC, intergenic, antisense, fusion, and genic isoforms combined (n = 27,002), 14,712 (54.5%) were classified as *bona fide* by SQANTI3. An additional 1,430 (5.3%) initially classified as artifacts were rescued based on multi-sample detection, read depth, and transcript-end support. The remaining 10,860 (40.2%) were excluded (**Methods, Supplementary Table 3**).

Overall, curation yielded 48,232 isoforms across 11,760 genes, retaining 77.0% of the 62,673 isoforms in the raw catalog. Retention varied across transcript categories, with 100% of FSM, 78.9% of NIC, 44.2% of NNC, and 41.1% of ISM isoforms retained (**Fig. 2a**).

**Figure 2.**
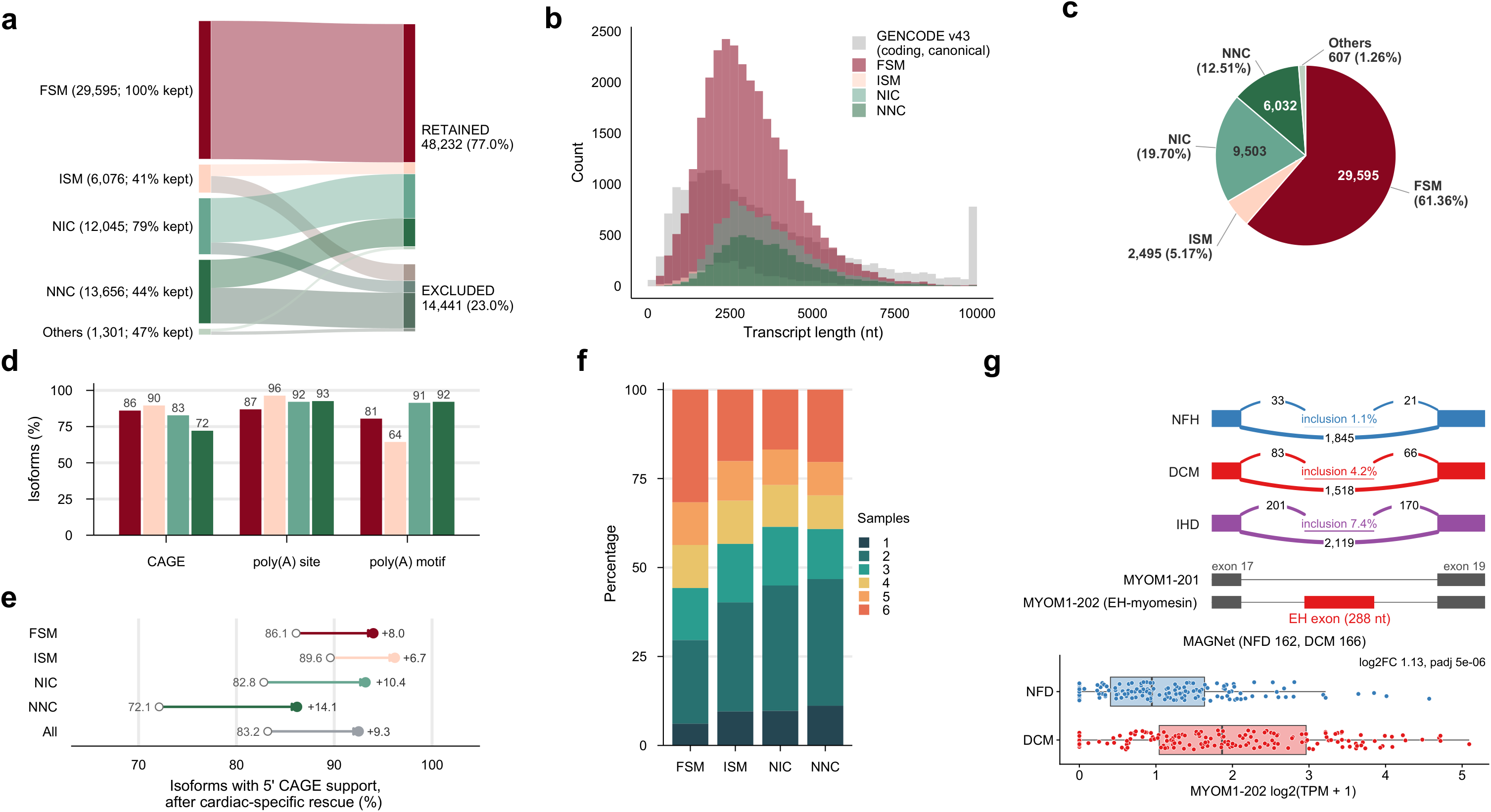
Curation and orthogonal validation of the cardiac isoform atlas. **a.** Sankey diagram illustrating the curation of detected isoforms into the final cardiac isoform atlas, stratified by transcript structural category. Numbers indicate the total number of detected isoforms and the percentage retained in the curated catalog: FSM (29,595; 100%), ISM (6,076; 41%), NIC (12,045; 79%), NNC (13,656; 44%), and other categories (1,301; 47%). **b.** Transcript-length distributions of curated isoforms across structural categories, compared with canonical protein-coding transcripts from GENCODE v43. **c.** Structural composition of the final curated atlas: FSM (61.4%), NIC (19.7%), NNC (12.5%), ISM (5.2%), and other transcript categories (1.3%). **d.** Orthogonal validation of transcript termini across structural categories, showing the proportion of isoforms supported by CAGE peaks, PolyASite 2.0 poly(A) sites, and poly(A) signal motifs. **e.** Improvement in 5′ end validation following cardiac-specific rescue of transcript start sites. Overall CAGE support increased from 83.2% to 92.5%, with the largest improvement observed for NNC isoforms (+14.1%). **f.** Distribution of curated isoforms across the six heart samples, showing the number of samples in which each isoform was detected, stratified by structural category. **g.** Long-read recovery of EH-myomesin, a known MYOM1 isoform reported to be re-expressed in dilated cardiomyopathy. Top, FLNC read coverage across MYOM1 exon 17-19, including the 288-nt EH exon, in NFH, DCM and IHD hearts (pooled reads). Bars show exon-level read counts relative to the larger flanking exon; upper and lower arcs indicate EH exon inclusion and skipping, respectively. Inclusion percentages (per heart): NFH, 1.7% and 0.3%; DCM, 6.4% and 1.8%; IHD, 1.2% and 14.3%. Reads were summed across curated isoforms. Middle, MYOM1-201 and MYOM1-202 (EH-myomesin), with the EH exon highlighted in red, introns are not to scale. Bottom, MYOM1-202 expression in MAGNet (NFH, n = 162; DCM, n = 166), shown as log2(TPM + 1). Boxes indicate the median and interquartile range; DESeq2 log2 fold change and adjusted p-value are for DCM versus NFH.

### Characterization and validation of the cardiac isoform reference

The curated reference had a mean isoform length of 3.26 kb (median 3.04 kb), comparable to GENCODE v43 isoforms (mean 3.65 kb, median 2.90 kb; **Fig. 2b**). FSM isoforms accounted for 61.4% of the reference, ISMs for 5.2%, and NIC + NNC isoforms for 32.2%, with fusion, genic, antisense, and intergenic isoforms comprising the remaining 1.3% (**Fig. 2c**).

Genes expressed a mean of 4.1 isoforms and a median of 2, with 67.3% (7,912) expressing more than one isoform. Isoform diversity was highly uneven across genes: 32.7% (3,848) expressed a single isoform, 14.5% (1,710) expressed 6-10 isoforms, and 175 genes (1.5%) expressed more than 20 (**Supplementary Figure 5).** NIC + NNC isoforms were distributed across 4,780 genes at a mean of 3.2 per gene. The most isoform-rich loci were dominated by large sarcomeric and cytoskeletal genes, including CAST (146 isoforms, 108 novel), SLMAP (86, 76 novel), LDB3 (64, 47 novel), and NEBL (47, 29 novel). At these loci, most detected transcripts were novel, highlighting substantial transcript diversity not represented in current annotations.

To assess the biological plausibility of identified isoforms, we cross-referenced transcript ends with orthogonal genomic resources: 5′ ends (TSSs) against a CAGE-peak reference^29^, and 3′ ends (TTSs) against PolyASite 2.0^30^ and a curated set of canonical and variant human poly(A) signal motifs^31,32^. FSM isoforms, which were retained without end filtering, showed strong support at both ends (CAGE 86.1%, PolyASite 86.9%, poly(A) motif 80.5%; **Fig. 2d, Supplementary Table 3**). ISMs showed similarly high CAGE (89.6%) and PolyASite (96.4%) support, consistent with the transcript-end criteria used during curation. Importantly, novel isoforms showed comparable or stronger 3′ end support despite end evidence being used to retain only a small minority of NIC + NNC isoforms (1,405 of 15,535). PolyASite and poly(A)-motif support exceeded 91% for both NIC and NNC isoforms, providing independent evidence for the authenticity of their 3′ ends. At the 5′ end, pan-tissue CAGE supported 82.8% of NIC and 72.1% of NNC isoforms. Since pan-tissue CAGE may underrepresent tissue-specific transcription start sites, we re-evaluated 5′ ends against a human cardiac CAGE atlas (55,204 robust promoter peaks; GSE150736^33^; **Methods**; **Supplementary Table 7**). The cardiac reference increased overall 5′-end support from 83.2% to 92.5%, with the largest gains observed for NNC (72.1% to 86.2%) and NIC (82.8% to 93.2%) (**Fig. 2e**).

Reproducibility across samples provided an additional line of support (**Fig. 2f**). Single-sample isoforms were uncommon across categories (FSM 6.1%, ISM 9.5%, NIC 9.7%, NNC 11.2%). FSM isoforms formed a robust core, with 70.4% detected in three or more samples and 31.7% detected in all six. Novel isoforms were less reproducible but remained substantial, with 55.1% of NIC and 53.3% of NNC detected in at least three samples and 16.8% and 20.3%, respectively, detected in all six. Because 1,405 novel isoforms were rescued partly based on detection in at least two samples, this criterion may have modestly increased their reproducibility rates.

The catalog also recovered known low-abundance cardiac isoforms. EH-myomesin, a MYOM1 isoform that includes a 288-nt EH exon and is re-expressed in dilated cardiomyopathy^34^, was detected as a full-splice match to MYOM1-202 (PB.10246.2). Pooled over the two hearts in each group, the EH exon was included in 1.1% of long reads spanning it in non-failing hearts, 4.2% in DCM and 7.4% in IHD, with variation between hearts (**Fig. 2g**), and MYOM1-202 was higher in DCM in MAGNet (log2FC = 1.13, FDR = 5e-6). Together, independent transcript-end support and cross-sample reproducibility provide orthogonal evidence for the reliability of the curated cardiac isoform reference.

### Independent RNA-seq support for novel splice junctions, exons, and transcript boundaries

We next evaluated transcript novelty using orthogonal short-read RNA-seq data from the MAGNet cohort (n = 366)^35^ and an independent dataset (GSE116250)^11^. Validation was stratified by the specific novel feature present in each isoform, including splice junctions, exonic regions, and transcript boundaries.

Of 9,014 novel isoforms containing at least one novel splice junction, 7,367 (81.7%) had strong support for all novel junctions in MAGNet (≥3 reads in ≥3 samples). An additional 1,166 isoforms increased the total to 8,533 (94.7%) with support for every novel junction in at least one sample. Only 481 (5.3%) lacked support for one or more novel junctions and were flagged for caution (**Fig. 3a**). Similar results were observed in the independent GSE116250 dataset, where 82.6% of novel isoforms had support for every novel junction despite the substantially smaller cohort size.

**Figure 3.**
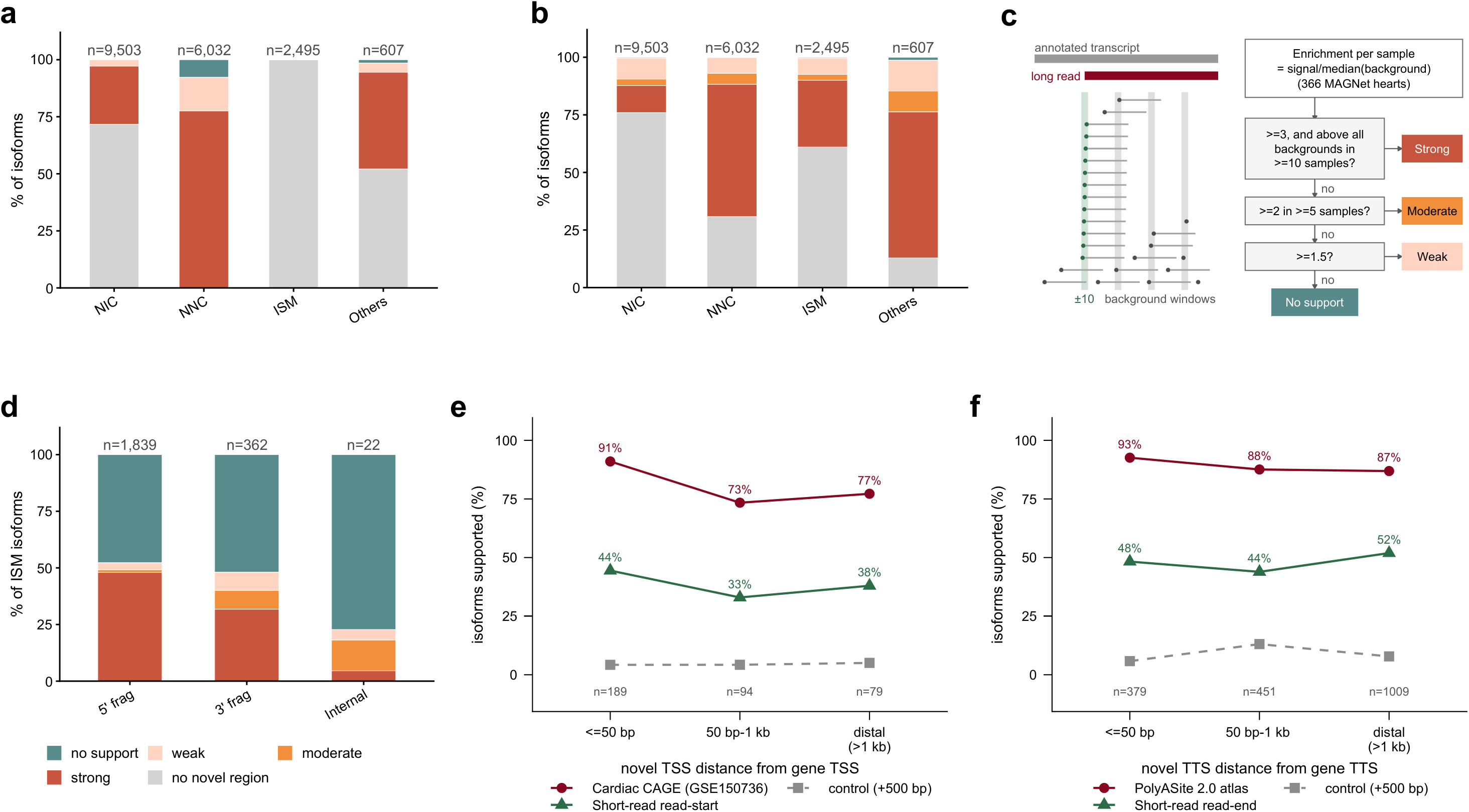
Independent validation of 18,637 non-FSM long-read isoforms using short-read evidence. **a, b, d**, Stacked bar plots showing the distribution of isoforms across levels of short-read support, with total isoforms counts indicated above each bar. Support ranges from no support or insufficient evidence (teal), through weak (pink) and moderate (orange), to strong (red). Light gray indicates test that was not applicable. **a.** Validation of splice-junction structures based on short-read evidence, distinguishing isoforms with all, some, or none of their junctions supported by short reads. **b.** Validation of novel transcript regions based on short-read coverage across the genomic regions underlying isoform novelty. **c.** Short-read validation of novel ISM transcript ends, illustrated for the novel 5′ end (TSS) of a 3′ fragment. Short-read 5′ ends are counted within a ±10-bp window at the candidate TSS and three background windows centered 100, 200, and 300 bp downstream. Enrichment is calculated per sample across 366 MAGNet hearts as the signal count divided by the median background count, and each end is classified according to the criteria shown. For novel 3′ ends of 5′ fragments, read 3′ ends are counted, with background windows positioned upstream. **d.** Validation of transcript ends for ISM isoforms, stratified by ISM subtype (5′ fragment, 3′ fragment, and internal fragment). **e.** Novel TSS support as a function of distance from the annotated gene TSS. Cardiac CAGE (GSE150736) support was 91% (<=50bp), 73% (50bp-1kb), and 77% (>1 kb); short-read support (strong or moderate enrichment) was 44%, 33%, and 38%, respectively. The gray dashed line shows the negative control using the same CAGE test with each candidate TSS shifted 500 bp into the gene body (4-5% across all distance bins). **f.** Novel TTS support as a function of distance from the annotated gene TTS. PolyASite 2.0 support was 93%, 88% and 87%; short-read support (strong or moderate enrichment) was 48%, 44% and 52%, respectively. The gray dashed line shows the negative control using the same PolyASite test with each candidate TTS shifted 500 bp into the gene body (6-13%).

Subtracting annotated exonic intervals identified 9,952 novel exonic regions (median length 80 bp) across 7,956 isoforms. Strand-specific coverage provided strong support for the novel exonic regions in 5,684 isoforms (71%), moderate support in 677 (9%), weak support in 1,535 (19%), and no support in 60 (<1%; **Fig. 3b, Methods**). Strongly supported regions were longer and exhibited near-complete coverage (median length 115 bp, coverage fraction 0.96), whereas weakly supported regions were shorter and sparsely covered (median length 16 bp, coverage fraction 0.19). These weakly supported regions were therefore more consistent with minor boundary shifts or terminal extensions than with large unsupported exons.

Although ISMs share annotated intron chains and were therefore excluded from the novel isoform set, their transcript boundaries may represent bona fide alternative transcription start or termination sites rather than truncation artifacts. We therefore evaluated independent short-read support for their novel termini.

Among 2,495 retained ISMs, 2,201 were terminal fragments, predominantly 5′ fragments (1,839 versus 362 3′ fragments). 5′ fragments already outnumbered 3′ fragments in the raw catalog (3,357 versus 1,488), consistent with lower coverage at the 3′ than at the 5′ end of full-length transcripts (**Supplementary Figure 3a**), and curation widened the gap because the novel 3′ ends of 5′ fragments fell in poly(A) sites more often (55%) than the novel 5′ ends of 3′ fragments fell in CAGE peaks (24%). Among 5′ fragments, which introduce novel 3′ termini, 882 (48%) showed strong short-read enrichment at the inferred 3′ end relative to gene-body background (**Fig. 3c**; **Methods**), consistent with alternative polyadenylation. Among 3′ fragments, which introduce novel 5′ termini, 115 (32%) showed strong enrichment at the inferred 5′ end, consistent with alternative transcription start sites (TSSs). Overall, 997 of 2,201 ISM fragments (45%) showed independent support for their novel terminal feature (**Fig. 3d, Supplementary Table 4**).

To determine whether validation was driven by proximity to annotated transcript boundaries, we stratified novel termini by distance to the nearest annotated TSSs or TTSs. End-enrichment remained largely stable across distance bins. For 3′ fragments, strong or moderate support for novel 5′ termini (40% overall) was comparable for sites located ≤50 bp (44%), 50 bp-1 kb (33%), and >1 kb (38%) from the nearest annotation (**Fig. 3e**). For 5′ fragments, strong or moderate support for novel 3′ termini (49% overall) was highest among sites located >1 kb from annotated transcript ends (52%) (**Fig. 3f**). These results provide evidence that many ISM termini represent bona fide alternative TSSs and TTSs, including sites located far from existing annotations, rather than technical truncation artifacts.

### Novel-isoform diversification concentrates in sarcomere and cardiomyopathy genes

Pooling across phenotypes, we ranked GENCODE-annotated genes by a reference-based diversification ratio, which measures the expansion of observed novel isoform diversity relative to the number of annotated isoforms in GENCODE v43 (**Methods**). By this metric, 109 genes showed more than a twofold expansion in isoform diversity (log2 ratio > 1), with the largest expansions observed in several sarcomeric and cytoskeletal genes, including MYH7, MYOZ2, SORBS1, FHOD3, TANC1, NRAP, MAP4, and TTN (Supplementary Table 5).

We next performed gene set enrichment analysis against MSigDB C5 Gene Ontology biological-process (GO) and Human Phenotype Ontology (HPO) gene sets^36-38^. To account for gene complexity, detectability, and expression, we residualized the diversification ratio on exon count, annotation depth, long-read depth (FLNC reads), and short-read expression before enrichment testing (**Methods**). Genes with greater isoform diversification were significantly enriched for cardiac and sarcomeric programs (**Fig. 4a, c)**. Enriched terms included myofibril assembly (NES = 2.15, FDR = 0.002), sarcomere organization (NES = 2.12, FDR = 0.004; **Fig. 4b**), and muscle cell development (NES = 2.05, FDR = 5e-4). These enrichments were essentially unchanged when canonical transcript and coding-sequence lengths were added to the model and remained significant among genes with canonical transcripts shorter than 6 kb, which most reads span in full. Among disease phenotypes, the most enriched term was asymmetric septal hypertrophy, a hallmark of hypertrophic cardiomyopathy (12 genes; NES = 2.24, FDR = 0.003) (**Fig. 4c, d**). The broader hypertrophic and dilated cardiomyopathy terms were not enriched after adjustment.

**Figure 4.**
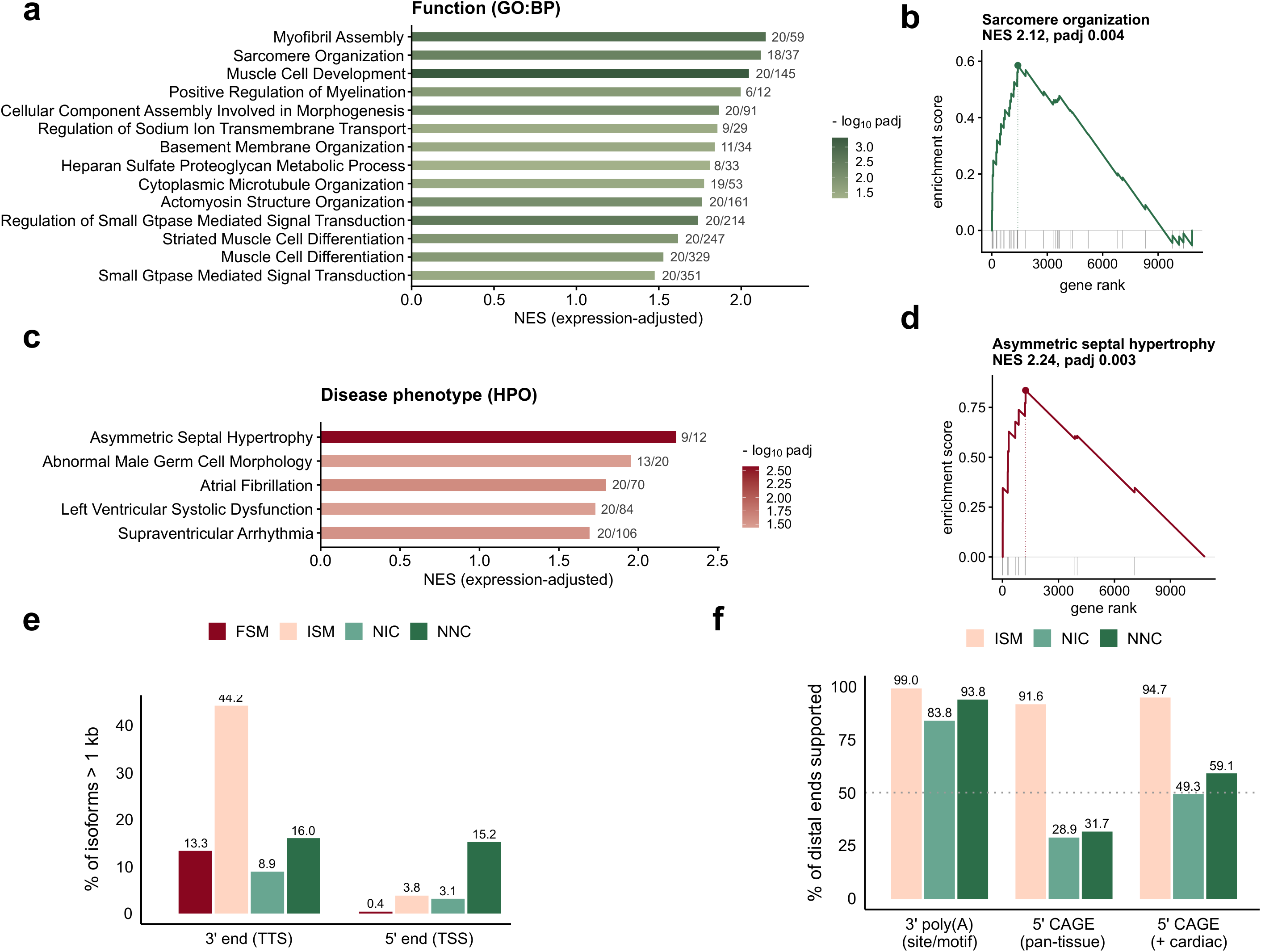
Functional enrichment and transcript-end diversification of cardiac genes. Gene-set enrichment analysis (GSEA) of genes ranked by reference-anchored novel-isoform diversification. Cardiac structural and disease related terms, including asymmetric septal hypertrophy, myofibril assembly, sarcomere organization and muscle-cell development, were significantly enriched, highlighting preferential diversification of sarcomere and cardiomyopathy-associated genes. **a.** Functional gene sets (GO:BP). Bars represent normalized enrichment score (NES), colored by -log10 adjusted p-value, with leading-edge/total set size annotated. **b.** Running enrichment score plot for sarcomere organization (NES = 2.12, adjusted p = 0.004), with genes ranked from highest to lowest diversification. **c.** Disease-phenotype gene sets (HPO). **d.** Running enrichment score plot for asymmetric septal hypertrophy (NES = 2.24, adjusted p = 0.003). **e.** Proportion of isoforms with distal transcript ends extending more than 1 kb beyond the reference annotation, shown separately for 3′ ends (TTS) and 5′ ends (TSS) by structural category. **f.** Proportion of distal transcript ends supported by independent evidence, including 3′ poly(A) sites/motifs, 5′ CAGE peaks from pan-tissue data, and 5′ CAGE peaks with cardiac-specific support, stratified by ISM, NIC, and NNC categories.

To confirm that these genes are expressed in heart muscle, we annotated them with single-cell expression from the Human Protein Atlas^39^. Of the 109 genes with more than twofold diversification, 100 were profiled in the atlas; 85 of these were expressed in cardiomyocytes (nCPM ≥ 20), and cardiomyocytes were the highest-expressing cardiac cell type for 58 (58% versus 19% of catalog genes; Fisher’s exact p = 1e-17). Genes with a long-read isoform differentially expressed in both short-read cohorts were also cardiomyocyte-dominant more often than catalog genes (40%; p = 9e-5).

Together, these results indicate that novel isoform diversification is disproportionately concentrated in sarcomeric and cardiomyopathy-associated genes, even after accounting for gene complexity, annotation depth, sequencing depth, and expression.

### NNC isoforms introduce novel boundaries and NMD substrates more frequently than NIC2

Of 15,535 novel isoforms in the final catalog, 9,503 were NIC and 6,032 were NNC. Compared with FSM, both NIC and NNC isoforms had higher exon counts (median 14 and 12 vs. 9; Mann-Whitney p < 1e-300 and p = 6.8e-199, respectively) and longer transcript lengths (median 3,349 bp and 3,452 bp vs. 2,887 bp; p = 3.7e-195 and p = 2.5e-197; **Supplementary Figure 6**).

NIC and NNC differed more clearly in transcript-boundary architecture. NNC isoforms more frequently introduced distal transcription start sites, with 15.2% of TSSs located >1 kb from annotated starts, compared with 3.1% for NIC and 0.4% for FSM (Fisher’s exact test, p < 1e-300 and p = 2.25e-92, respectively). NNC isoforms also showed more distal transcription termination sites, with 16.0% of TTSs located >1 kb from annotated ends, compared with 13.3% for FSM and 8.9% for NIC (p = 2.57e-8 and p = 4.03e-32, respectively; **Fig. 4e**). For example, the NNC isoform PB.1169.9 of OBSCN starts 2.8 kb from the nearest annotated transcript start and 161 kb downstream of the canonical OBSCN promoter, within a cardiac CAGE cluster (3,279 tags), and the splice junction of its unannotated first exon was detected in 204 of 366 MAGNet hearts (**Supplementary Figure 7**). Thus, NIC isoforms more often retained annotated transcript boundaries, whereas NNC isoforms more frequently combined novel splice structures with distal transcript boundaries.

Among isoforms with TTSs located >1 kb from annotated ends, 93.8% of NNC and 83.8% of NIC isoforms overlapped a poly(A) site or motif. Distal TSSs showed weaker support in the pan-tissue CAGE reference, with overlap for 31.7% of NNC and 28.9% of NIC isoforms. Re-evaluation using the human cardiac CAGE atlas increased support to 59.1% and 49.3%, respectively (**Fig. 4f**; **Supplementary Table 7**). Thus, most distal TTSs and a substantial fraction of distal TSSs have independent transcript-end support.

Finally, we assessed the potential functional consequences of these structural differences by predicting open reading frames and nonsense-mediated decay (NMD) susceptibility^24,40^. Most NIC and NNC isoforms were predicted to be protein-coding (98.1% and 96.2%, respectively), similar to FSM isoforms (96.3%). However, NNC isoforms were substantially more likely to contain predicted NMD-sensitive premature termination codons (12.9%) than NIC (4.9%) or FSM (1.1%) isoforms (**Supplementary Figure 6c**). These results indicate that NNC isoforms are not only structurally more divergent from annotated transcripts but are also more likely to generate transcripts subject to post-transcriptional NMD.

### Population-scale expression of the long-read isoform catalog

Given the modest sample size of our long-read cohort, we next evaluated expression of the 18,637 non-FSM isoforms (NIC, NNC, ISM, and minor categories) in the independent MAGNet cohort of 366 left-ventricular samples profiled by short-read RNA-seq. Short reads were quantified against the long-read transcript reference using a merged-reference Salmon^41^ decoy pipeline^42^ (**Fig. 5a**; **Methods**).

**Figure 5.**
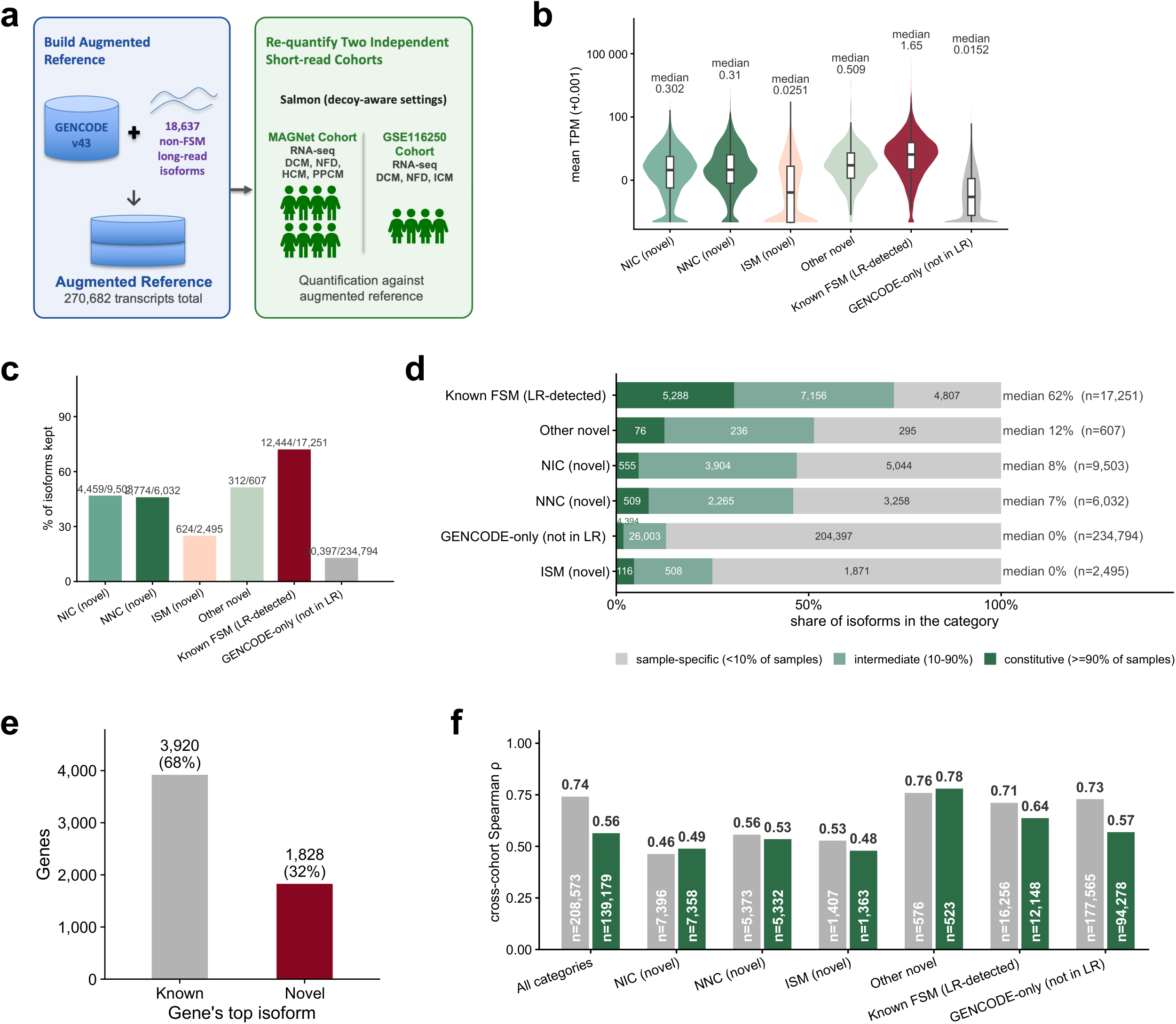
Population-scale expression and cross-cohort reproducibility of novel isoforms. **a.** Short-read quantification workflow. GENCODE v43 was combined with the 18,637 non-FSM long-read isoforms to generate a reference containing 270,682 transcripts. MAGNet (n = 366; DCM, NFH, HCM, and PPCM) and GSE116250 cohorts were re-quantified using Salmon with decoy-aware settings. Panels b-f use these expression estimates. **b.** Mean transcript expression (TPM) by category; medians expression was lowest for ISM and GENCODE-only transcripts and highest for known FSM isoforms. **c.** Short-read detection rates by category, with the number of detected isoforms indicated above each bar. **d.** Distribution of isoform detection breadth across samples, classified as sample-specific, intermediate, or constitutive. Median detection (TPM >= 1) was 62% of samples for known FSM isoforms, compared with 0-12% for non-FSM categories. **e**. Classification of genes according to whether their most highly expressed isoform was known (3,920; 68%) or novel (1,828; 32%). **f.** Cross-cohort reproducibility of isoform expression, measured by Spearman correlation (ρ) between mean isoform TPM in MAGNet and GSE116250, among isoforms expressed in both cohorts, before and after abundance adjustment. Gray bars show unadjusted correlations; green bars show correlations after inverse-probability weighting to match each category to the abundance distribution of the pooled novel isoforms. This adjustment reduces the influence of differences in abundance distributions across categories on correlation estimates. The apparent advantage of GENCODE-only isoforms, which span the widest abundance range, was attenuated after adjustment (ρ = 0.73 to 0.57), bringing their correlation closer to those of NIC, NNC, and ISM isoforms (ρ = 0.48-0.53). Values above bars indicate ρ; numbers inside bars indicate the number of isoforms (gray) and effective sample size after weighting (green).

Non-FSM isoforms were expressed across the population but generally at lower abundance than FSM isoforms. Long-read-detected FSM isoforms had the highest median abundance (1.65 TPM), whereas NIC, NNC, and ISM isoforms had median abundances of 0.30, 0.31, and 0.025 TPM, respectively, all exceeding the GENCODE-only background (0.015 TPM) (**Fig. 5b**). Using a low-expression threshold of TPM ≥1 in ≥10% of samples, 43.8% of non-FSM isoforms (8,169/18,637) met this criterion, including 47% of NIC, 46% of NNC, and 51% of other non-FSM isoforms, compared with 72% of FSM isoforms and 13% of GENCODE-only isoforms (**Fig. 5c**). ISMs were substantially less prevalent, with 624 of 2,495 (25%) meeting the same threshold. Consistent with their lower abundance, non-FSM isoforms were detected in fewer samples overall than FSM isoforms, but their detection breadth remained substantially greater than that of GENCODE-only isoforms (**Fig. 5d**), indicating that these transcripts were not confined to a small number of isolated samples.

Although non-FSM isoforms were less abundant and detected in fewer samples than known isoforms (**Fig. 5d**), collectively they often accounted for a substantial share of gene expression in MAGNet (**Fig. 5e**). Among 5,358 annotated genes with at least one non-FSM isoform, non-FSM isoforms contributed a median of 18% of total gene TPM. Their contribution exceeded 50% in 1,177 genes (22.0%), and a non-FSM isoform was the most highly expressed transcript in 1,438 genes (26.8%). These dominant non-FSM isoforms were broadly expressed, reaching at least 1 TPM in a median of 90% of samples, compared with 93% for known top isoforms. Thus, the lower overall detection of non-FSM isoforms as a class largely reflects the many low-abundance isoforms rather than the expression patterns of the dominant ones.

We next assessed reproducibility in an independent cohort of 63 left-ventricular samples from GSE116250. Mean isoform expression showed strong concordance between MAGNet and GSE116250 across 208,573 isoforms (Spearman ρ = 0.74), and moderate concordance for non-FSM isoforms (ρ = 0.51). By isoform class, correlations were 0.56 for NNC, 0.53 for ISM, 0.46 for NIC, and 0.76 for other non-FSM isoforms, compared with 0.73 for GENCODE-only transcripts (**Fig. 5f**). Because cross-cohort correlations can be influenced by differences in abundance distributions, we reweighted each class to a shared abundance distribution; the GENCODE-only correlation then fell to 0.57, close to those of NIC (0.49), NNC (0.54), and ISM (0.48) (**Fig. 5f**). Down-sampling each class to matched abundance and size gave the same result: correlations for novel isoforms were comparable to those for GENCODE-only transcripts, with overlapping resampling intervals (**Supplementary Figure 8**).

Together, these analyses demonstrate that a substantial fraction of long-read-derived non-FSM isoforms is detectable in population-scale short-read data. Although some isoforms are less abundant and less broadly expressed than annotated FSM isoforms, many contribute substantially to gene-level expression and show reproducible expression patterns across independent cardiac cohorts.

### Novel coding isoforms remodel protein domain architecture

Among novel coding isoforms (NIC+NNC) whose gene has a protein-coding canonical GENCODE v43 transcript and in which either protein carries at least one Pfam domain (n = 14,747), 3,670 (24.9%) showed altered Pfam-domain content^43^ relative to the corresponding canonical GENCODE v43 protein, predominantly through domain loss (3,393 loss-only isoforms; **Fig. 6a**). These losses were terminal (39.8%), resulted from divergent reading frames over retained loci (35.6%), internal exon excision (16.4%), or sub-threshold domain erosion (8.2%), whereas domain losses in ISMs were almost entirely terminal (95.3% of 1,186; **Fig. 6b**). Frameshift-divergent losses were more frequently predicted to undergo NMD than internal exon-excision events (23.1% versus 4.1%), suggesting that some domain-loss transcripts may represent degradation substrates rather than stable protein products.

**Figure 6.**
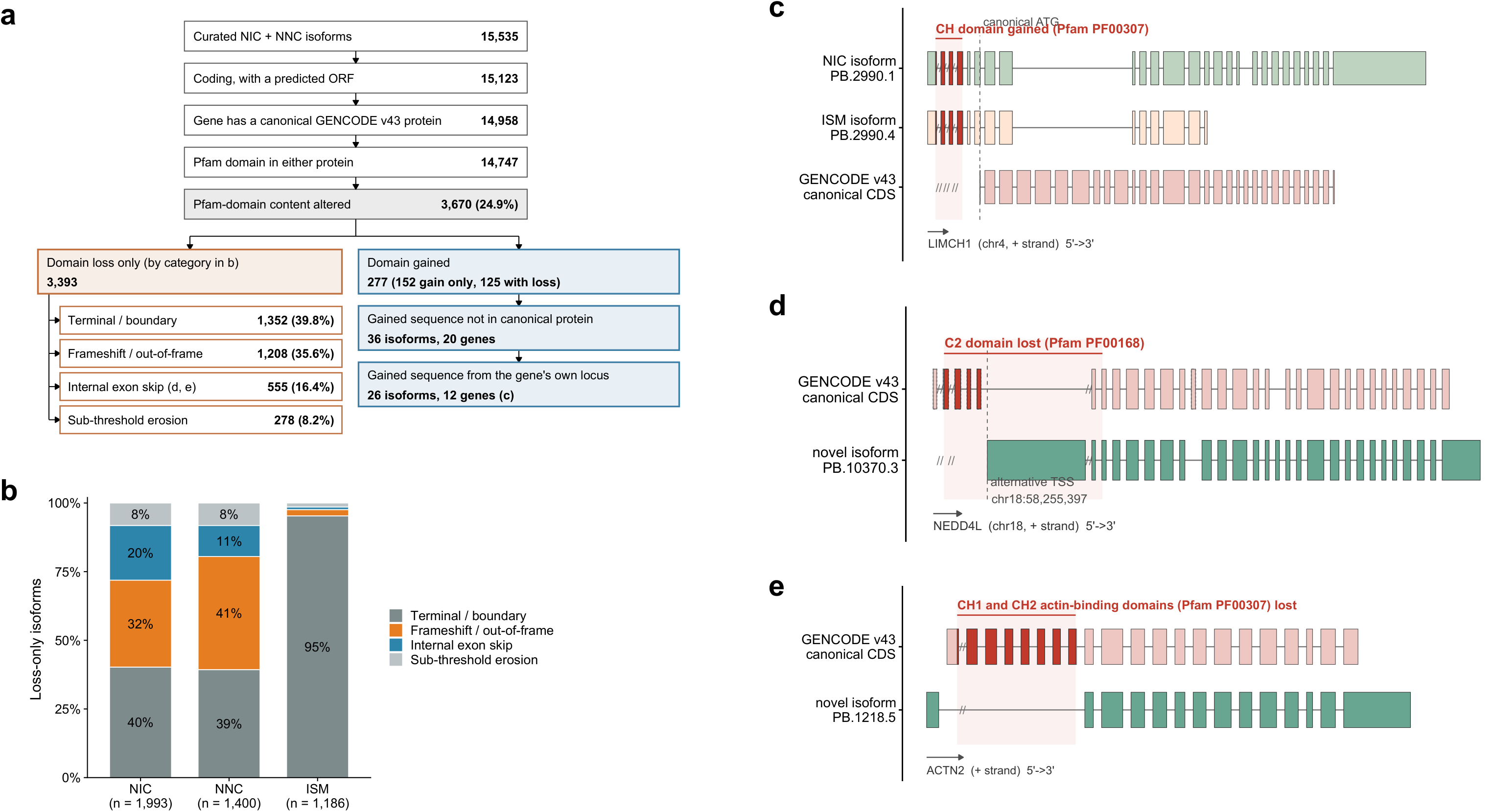
Predicted protein-domain consequences of novel isoforms. **a.** Comparison of predicted Pfam-domain content between curated coding NIC and NNC isoforms and the canonical GENCODE v43 protein encoded by the corresponding gene. Boxes indicate the number of isoforms retained at each step. Domain losses are classified into four mechanisms and pooled across NIC and NNC categories (category-specific results in b). Predicted domain gains are retained only when the gained sequence is absent from the canonical protein and originates from the gene’s own locus, excluding chimeric, read-through, and overlapping-gene sequences. Letters indicate the examples shown in c-e. **b.** Mechanisms of domain loss among loss-only isoforms in each category: terminal or boundary, frameshift or out-of-frame, internal exon skip, or sub-threshold erosion. **c.** LIMCH1 example: NIC isoform PB.2990.1 gains a CH domain (Pfam PF00307) through alternative TSS usage, compared with ISM isoform PB.2990.4 and the canonical GENCODE protein. **d.** NEDD4L example: novel isoform PB.10370.3 loses most of the C2 domain (Pfam PF00168) through initiation from an alternative first exon (chr18:58,255,397). **e.** ACTN2 example: novel isoform PB.1218.5, an N-terminally truncated isoform predicted to initiate translation at Met283, lacks both calponin-homology domains (Pfam PF00307) that comprise the actin-binding domain.

Apparent domain gains were largely attributable to differences in HMMER scoring^44^ between proteins of different lengths; requiring gained domain sequences to be poorly represented in the canonical protein and excluding chimeric and read-through transcripts reduced 277 initial candidates to 26 isoforms across 12 genes (**Fig. 6a**). In LIMCH1, for example, a 5′-extended ORF introduced an N-terminal calponin-homology (CH) actin-binding domain (Pfam PF00307) that was absent from the canonical protein (**Fig. 6c**).

Internal exon skipping removed N-terminal domains in NEDD4L and ACTN2 (**Fig. 6a**). In NEDD4L (**Fig. 6d**), a HECT-family E3 ubiquitin ligase that regulates the cardiac sodium channel NaV1.5^45,46^, fourteen isoforms initiated at an alternative first exon within canonical intron 5 (shared with the annotated transcript NEDD4L-238) and excluded the five 5′-most coding exons, which encode most of the N-terminal C2 domain. These isoforms replace the C2 region with a 378-residue N-terminal segment without known domains and retain one to three of the four WW domains and, in thirteen cases, the complete catalytic HECT domain. Similarly, ACTN2 isoform PB.1218.5 splices from the exon 1 5′ UTR directly to exon 9; translation from Met283 yields an N-terminally truncated α-actinin-2 that lacks both calponin-homology domains (Pfam PF00307) of the actin-binding domain^47^ (**Fig. 6e**).

Frameshift-dependent domain loss represented the second most common mechanism. APLN long-read isoform PB.12263.1 encodes a 100-amino-acid ORF that was not predicted to be NMD-sensitive but diverges from the canonical preproapelin reading frame and lacks the C-terminal Apelin domain (Pfam PF15360), which contains the bioactive peptide^48,49^ (**Supplementary Figure 9**).

Strict in-frame cassette excision defined a smaller class: 99 isoforms across 65 genes altered a protein domain while preserving the surrounding reading frame, including SMYD1, which recurrently lost its zf-MYND module, and GIT2 isoform PB.7846.10, which removed 50 residues of its GIT_CC coiled-coil domain. Overall, domain losses far outnumbered gains, consistent with previous reports that alternative splicing more frequently removes than creates folded protein domains^50,51^.

### Augmented quantification captures disease-regulated isoforms in independent short-read cohorts

We next asked whether the expanded transcript catalog captures disease-associated isoform regulation by testing both cohorts’ transcript-level estimates (**Fig. 5a**) for differential expression between diseased and non-failing hearts.

In the MAGNet cohort, among covariate-complete samples (n = 328; DCM = 166, NFH = 162), 76,021 transcripts passed expression filtering. Differential expression analysis comparing DCM with NFH using DESeq2^52^, with adjustment for age, race and sex (**Fig. 7a**), identified 24,081 differentially expressed transcripts at FDR < 0.05, including 3,356 with more than 2-fold changes (1,428 up-and 1,928 down-regulated in DCM). Of the 18,637 non-FSM isoforms, 11,761 (63%) passed filtering, and 594 showed more than 2-fold changes (153 up- and 441 down-regulated), accounting for 17.7% of all transcripts with more than 2-fold changes. This demonstrates that the long-read atlas captures disease-regulated transcripts beyond those represented in standard GENCODE annotations. Most of these 594 transcripts (469, 79%) contained a novel junction, novel exonic sequence, or distinguishing exon pair directly observable by short reads. The remaining 125 differed from annotated transcripts only through distant exon combinations or transcript ends, making their short-read estimates dependent on read assignment (**Methods**).

**Figure 7.**
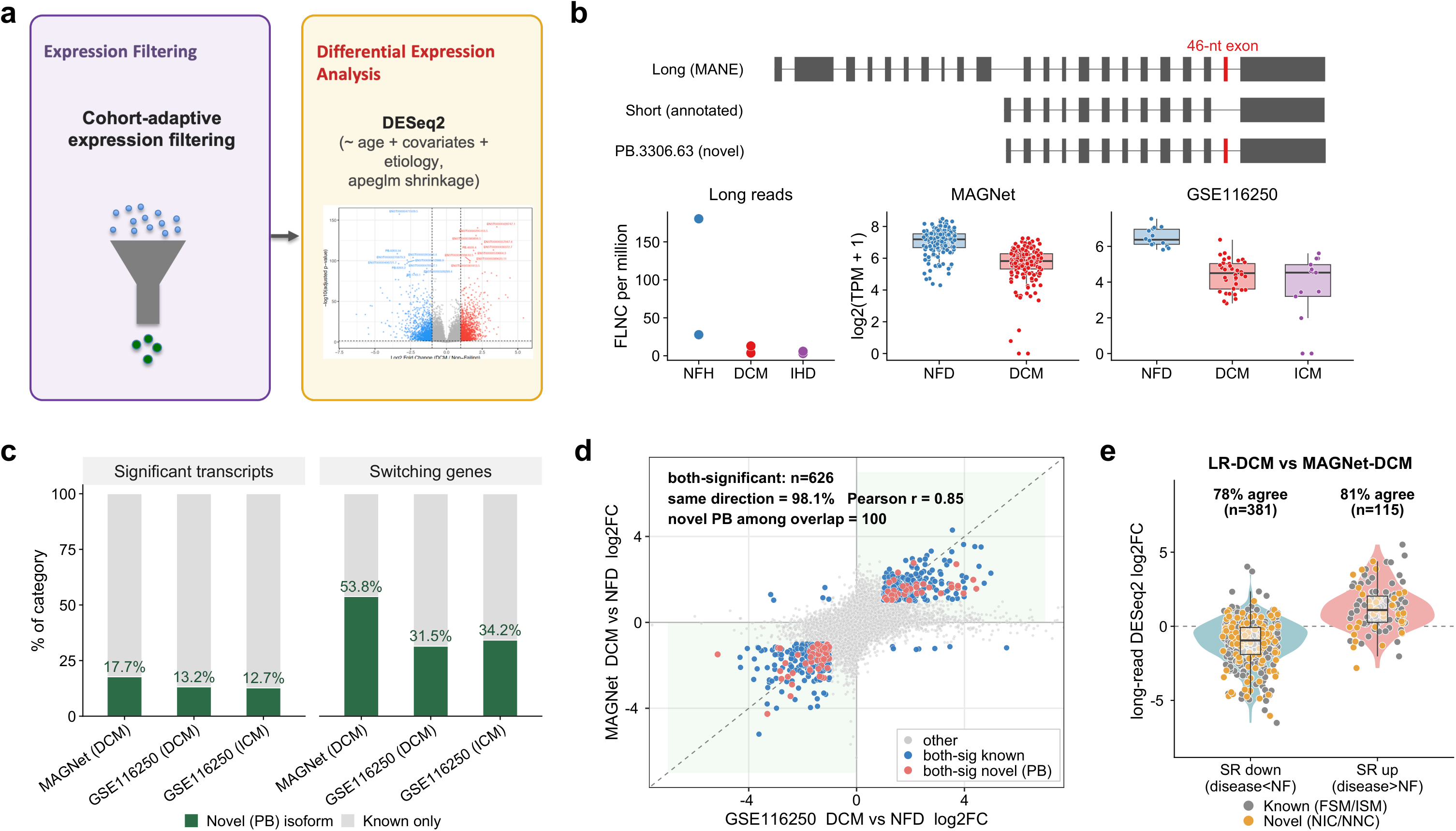
Novel isoforms contribute to disease-associated expression changes. **a.** Differential-expression analysis workflow, including cohort-adaptive filtering of the Salmon expression estimates described in Fig. 5a, followed by DESeq2 analysis with covariate adjustment and apeglm log2 fold-change shrinkage. **b.** Example of a novel short isoform of PALLD (PB.3306.63). Top, canonical long isoform (MANE, ENST00000505667), annotated short isoform (ENST00000704822), and PB.3306.63, which uses the promoter of the annotated short isoform but retains the 46-nt exon of the canonical isoform (red); introns are compressed. Bottom, PB.3306.63 expression in long-read data (FLNC reads per million retained FLNC reads; one point per heart), MAGNet (NFH, n = 162; DCM, n = 166), and GSE116250 (NFH, n = 14; DCM, n = 36; ICM, n = 13), shown as log2(TPM + 1). Boxes indicate the median and interquartile range. **c.** Contribution of non-FSM isoforms to disease-associated expression signals across MAGNet-DCM, GSE116250-DCM, and GSE116250-ICM. **d.** Cross-cohort comparison of transcript-level differential expression for DCM versus NFH, independently estimated in MAGNet (y-axis) and GSE116250 (x-axis) using the same augmented reference. Each point represents a transcript quantified in both cohorts. Gray points indicate all co-tested transcripts; blue points indicate transcripts significant in both cohorts (adjusted p < 0.05 and |log2FC| > 1); red points highlight the non-FSM long-read isoforms among these shared significant transcripts. The dashed line indicates the identity line (y = x), and shaded green quadrants indicate concordant directions of change. **e.** Directional concordance of DCM-associated expression changes between the long-read cohort and MAGNet. Points represent 496 long-read isoforms that met the expression threshold for long-read differential-expression testing (at least 10 FLNC reads in total and detected in at least two of six hearts) and whose corresponding MAGNet transcripts were differentially expressed in DCM versus NFH (adjusted p < 0.05 and |log2FC| > 1). Each point shows the long-read DESeq2 log2FC (DCM versus NFH), grouped by the direction of the corresponding MAGNet change (down, teal; up, red). Novel isoforms (NIC/NNC, gold) were matched to MAGNet by their own identifiers, whereas known isoforms (FSM/ISM, gray) were matched by their associated GENCODE transcripts; accordingly, 349 known isoforms correspond to 242 MAGNet transcripts. Box plots indicate the median and interquartile range; the dashed line marks zero.

An example was PB.3306.63, a novel short isoform of PALLD (palladin) that accounted for 57% of PALLD expression in NFHs. It starts at the internal promoter of the annotated short isoform but includes a 46nt exon from the canonical isoform, thereby encoding the canonical C-terminus with its three C-terminal Ig domains (**Fig. 7b**). PB.3306.63 was reduced in DCM in MAGNet (log_2_FC = -1.51) and in GSE116250 (DCM, -2.18; ICM, - 2.02), consistent with the long-read data (85 FLNC reads in the two NFHs versus 5 and 3 in the DCM and IHD pairs).

Among the differentially expressed transcripts, 88 high-confidence transcripts mapped to 26 genes containing both up- and down-regulated isoforms, consistent with isoform switching. These included SORBS2, ANXA11, FHL1, NRAP, and SNHG14, with within-gene log_2_FC ranges of up to 7.5. Fourteen of the 26 genes (54%) involved a long-read isoform (**Fig. 7c**). a formal test of differential isoform usage using IsoformSwitchAnalyzeR with satuRn^53,54^ identified significant switches in 39 genes.

We next evaluated the independent GSE116250 cohort, which included DCM (n = 36), ICM (n = 13), and NFH (n = 14) samples. Differential expression analysis identified 4,203 significant transcripts (FDR < 0.05, more than 2-fold change) in DCM versus NFH and 9,154 in ICM versus NFH. Genes containing both up- and down-regulated transcripts numbered 197 in DCM and 928 in ICM, with 151 of the 197 DCM genes (77%) also identified in ICM. Long-read isoforms accounted for 555 of the 4,203 significant transcripts in DCM (13.2%) and 1,166 of the 9,154 significant transcripts in ICM (12.7%). They also contributed to isoform switching in 62 of 197 DCM genes (31.5%) and 317 of 928 ICM genes (34.2%), indicating that approximately one-third of isoform-switching genes involved transcripts not represented in standard reference annotations (**Fig. 7c**).

To test whether these transcript-level changes were detectable at the gene level, we repeated the analysis using gene-level counts. Of the 2,311 MAGNet genes with a differentially expressed transcript (FDR < 0.05, more than 2-fold), 1,334 (58%) showed less than 2-fold changes at the gene level, including 330 of 435 genes (76%) with differentially expressed long-read isoform and 25 of 26 isoform-switching genes (96%). Similar patterns were observed in GSE116250 (DCM, 78% and 84%; ICM, 84% and 89%; **Supplementary Table 6**), indicating that gene-level analysis obscures much of the disease-associated transcript remodeling.

To assess the reproducibility of DCM-associated isoform changes, we compared transcript-level DCM-versus-NFH log2FC values between MAGNet and GSE116250. Across 51,741 transcripts testable in both cohorts, including 7,359 long-read isoforms, fold changes were positively correlated (Pearson r = 0.30; Spearman ρ = 0.43). Concordance increased substantially among the 626 transcripts that were differentially expressed in both cohorts (r = 0.85), with 98.1% changing in the same direction (**Fig. 7d**). Notably, 100 of these 626 reproducibly differential transcripts were long-read only isoforms, and all 100 changed in the same direction across cohorts, compared with 97.7% of annotated transcripts. Of the 100 long-read isoforms, 86 carried a novel feature directly observable by short reads. Among all MAGNet differentially expressed transcripts testable in GSE116250, 89% changed in the same direction. Thus, the expanded long-read reference identifies disease-associated isoform changes that are reproducible across independent patient cohorts.

Finally, we examined whether isoform-switching patterns differed between DCM and ICM in GSE116250. The two diseases showed broadly similar, but not identical, isoform-switching patterns. Measured against separate halves of the NFH group, changes in DCM and ICM were correlated (r = 0.64), comparable to the correlation between two separate groups of 13 DCM hearts (r = 0.65). All 37 isoforms that switched significantly in either disease changed in the same direction in the other. A direct comparison of ICM with DCM identified only 10 differing isoforms across 8 genes, 9 of which did not switch significantly relative to NFH in either disease (**Supplementary Fig. 10**).

### Cross-platform validation of long-read-derived disease-associated isoforms

We next assessed whether disease-associated isoform changes identified in the long-read cohort were reproducible in independent short-read cohorts. We compared each disease phenotype with NFH using per-sample FLNC counts for 17,945 robustly detected long-read isoforms (**Methods**). Given the limited sample size of the long-read cohort, cross-cohort reproducibility was assessed based on the direction of expression change rather than fold-change magnitude. Among the 3,356 isoforms confidently differential in MAGNet (FDR < 0.05, |log2FC| > 1), 496 were sufficiently expressed for evaluation in the long-read data, including 349 known (FSM/ISM) and 147 novel (NIC/NNC) isoforms. Long-read and MAGNet fold changes agreed in direction for 78.6% of isoforms (binomial p = 1.7e-39; **Fig. 7e**), with similar concordance for novel isoforms (78.9%). In the independent GSE116250 cohort, directional concordance was lower (58.6% of 645 isoforms; p = 7e-6; **Supplementary Figure 11**) but remained above chance. Of the 17 isoforms significant by long-read omnibus likelihood-ratio testing (**Supplementary Figure 12**), 14 were testable in MAGNet: 11 were significantly changed in the same direction (representing nine distinct MAGNet transcripts), and three showed concordant but nonsignificant changes. The remaining three were insufficiently expressed in MAGNet. In contrast, directional reproducibility was limited for ischemic disease. Long-read IHD fold changes agreed with those from GSE116250 ischemic cardiomyopathy for 53.6% of 1,386 confidently differential isoforms (**Supplementary Figure 11**), only marginally above chance (one-sided binomial p = 0.004). Thus, DCM-associated isoform changes showed directional concordance with larger independent cohorts, whereas the IHD signal did not. Given the near-identical DCM and ICM isoform changes in GSE116250, this discrepancy likely reflects the two IHD hearts sampled rather than a distinct ischemic splicing program.

### CardioIsoMap provides a long-read isoform resource for heart research

We developed CardioIsoMap, a web portal providing access to 48,232 curated isoforms across 11,760 genes identified by long-read sequencing of failing and non-failing human hearts (https://cardioisomap.streamlit.app/; **Fig. 1a**). The portal integrates isoform structures with functional annotations, including isoform class, coding potential, predicted NMD status, transcript-end support (CAGE peaks and poly(A) motifs) and, for novel isoforms, splice-junction support from short-read RNA-seq. Users can explore isoform distributions globally or at individual genes, visualize transcript structures alongside GENCODE reference models, and compare long-read isoform usage across phenotypes with short-read isoform expression in the independent MAGNet cohort.

## Discussion

Reference annotations of the human transcriptome are derived largely from healthy and non-cardiac tissues, whereas short-read sequencing reconstructs transcripts indirectly. Consequently, cardiac isoform diversity, particularly in disease, remains incompletely characterized. By sequencing full-length transcripts from failing and non-failing human left ventricle, we quantified unannotated cardiac isoform diversity and evaluated its reproducibility and biological support. Our results reveal a substantial and structured layer of cardiac transcript variation that is absent from current annotations and is supported across independent datasets and sequencing platforms.

A central finding is that observed isoform diversity is strongly influenced by sampling depth. The relationship between gene expression and isoform count was substantially reduced after controlling for sequencing depth, indicating that raw isoform counts partly reflect sampling effort rather than intrinsic transcript complexity. This has important implications for interpreting isoform diversity across genes, particularly when sequencing depth varies. After accounting for depth, exon count was primarily associated with splice diversity, whereas transcript length was more strongly associated with variation at transcript termini. Thus, controlling for sampling depth is essential when interpreting isoform diversity as a biological property.

Multiple lines of evidence support the biological validity of the novel isoforms identified here. Novel splice junctions were supported by independent short-read datasets, while transcript boundaries showed evidence consistent with alternative transcription initiation and termination. Novel isoforms were enriched in genes encoding sarcomeric and cytoskeletal proteins, and these associations persisted after adjustment for expression, transcript length, annotation depth, and sequencing depth. At the protein level, novel coding isoforms predominantly altered existing protein architectures through truncation, domain loss, or changes in transcript termini rather than introducing new domains. In-frame internal exon excisions provided additional examples of transcript variation that can preserve the reading frame while altering functional domain content, suggesting a potential mechanism for protein-level diversification.

Isoform-level analysis further revealed disease-associated transcriptional changes that were not apparent from gene-level expression alone. Many genes showed changes in isoform usage without large changes in overall gene expression, indicating that cardiac disease-associated remodeling can occur at the transcript level. A substantial fraction of these events involved transcripts absent from standard annotations, highlighting how incomplete transcript models can obscure aspects of cardiac disease biology. Genes with disease-associated isoform changes included sarcomeric and cytoskeletal genes such as PALLD, SORBS2, FHL1, and NRAP.

In the short-read cohort that included both etiologies, isoform changes in DCM and ICM were nearly identical: their agreement reached the level expected between two separate groups of DCM hearts, and every isoform that switched significantly in one disease changed in the same direction in the other. This agrees with a recent long-read study of 33 left ventricles in which DCM and ICM isoform landscapes were largely indistinguishable^22^, and suggests that end-stage heart failure converges on a shared isoform program regardless of etiology. The weak replication of the long-read IHD signal therefore more likely reflects the two IHD hearts we sequenced than a distinct ischemic program, and the few switches that differed directly between ICM and DCM need confirmation in larger cohorts.

These findings extend previous long-read studies in cell lines, brain, and heart tissue^16-18,21,22^ that have demonstrated extensive transcript diversity beyond existing annotations. Here, we show that much of this novel cardiac isoform content is supported by independent evidence and can be quantified and replicated at population scale, and further demonstrate the complementary value of long- and short-read sequencing. Long-read sequencing provides direct resolution of transcript structures, whereas larger short-read cohorts provide greater statistical power for population-scale expression analysis and disease association. Integrating these complementary strengths therefore provides a practical framework for characterizing transcript diversity and evaluating disease-associated isoform changes.

To facilitate exploration of this resource, we developed CardioIsoMap, an isoform-centric web portal integrating transcript structure, annotation, and expression. CardioIsoMap enables direct interrogation of isoform-level features and disease-associated expression patterns and provides a resource for hypothesis generation and downstream functional studies.

Several limitations should be considered. The long-read discovery cohort was small and was not powered for standalone differential expression analysis, motivating validation in larger short-read cohorts. Isoform discovery did not reach saturation, indicating that the present catalog likely represents a lower bound on cardiac transcript diversity. End-coverage bias, particularly for long transcripts, may contribute to ISM annotations and to apparent novel isoforms at long genes such as TTN. Bulk tissue profiling also precludes cell-type resolution, which will require single-cell or single-nucleus long-read approaches. Finally, predicted coding consequences, NMD sensitivity, and protein-domain changes remain computational inferences and require experimental validation.

In summary, full-length sequencing of the human heart reveals a substantial and structured layer of isoform diversity not captured by current annotations. Proper interpretation of this diversity requires accounting for sampling depth, while disease-associated remodeling is frequently evident at the isoform level even when gene-level expression changes are modest. Replicated DCM-associated isoform changes and protein-domain remodeling events provide a foundation for future mechanistic studies. Larger cohorts, cell-type-resolved profiling, and experimental validation will be needed to establish the functional and disease relevance of these transcriptomic changes.

## Acknowledgments

We thank the investigators of the Myocardial Applied Genomics Network (MAGNet; GSE141910), of GSE116250, and of the human cardiac CAGE atlas (GSE150736) for making their data publicly available. Computational analyses were performed on the high-performance computing cluster of GENIE, University of Texas Health at San Antonio. This work was supported by the National Heart, Lung, and Blood Institute of the National Institutes of Health under award number R01HL153403 (F.L.) and 1R21HL184221-01 (X.W.). Claude and ChatGPT were used for language editing and improving the clarity and readability of the manuscript.

## References

1 Roth, G. A. et al. Global Burden of Cardiovascular Diseases and Risk Factors, 1990-2019: Update From the GBD 2019 Study. J Am Coll Cardiol 76, 2982–3021 (2020). 10.1016/j.jacc.2020.11.010

2 McNally, E. M. & Mestroni, L. Dilated Cardiomyopathy: Genetic Determinants and Mechanisms. Circ Res 121, 731–748 (2017). 10.1161/CIRCRESAHA.116.309396

3 Weintraub, R. G., Semsarian, C. & Macdonald, P. Dilated cardiomyopathy. Lancet 390, 400–414 (2017). 10.1016/S0140-6736(16)31713-5

4 Frangogiannis, N. G. The inflammatory response in myocardial injury, repair, and remodelling. Nat Rev Cardiol 11, 255–265 (2014). 10.1038/nrcardio.2014.28

5 Wang, E. T. et al. Alternative isoform regulation in human tissue transcriptomes. Nature 456, 470–476 (2008). 10.1038/nature07509

6 Reyes, A. & Huber, W. Alternative start and termination sites of transcription drive most transcript isoform differences across human tissues. Nucleic Acids Res 46, 582–592 (2018). 10.1093/nar/gkx1165

7 van den Hoogenhof, M. M., Pinto, Y. M. & Creemers, E. E. RNA Splicing: Regulation and Dysregulation in the Heart. Circ Res 118, 454–468 (2016). 10.1161/CIRCRESAHA.115.307872

8 Guo, W. et al. RBM20, a gene for hereditary cardiomyopathy, regulates titin splicing. Nat Med 18, 766–773 (2012). 10.1038/nm.2693

9 Kong, S. W. et al. Heart failure-associated changes in RNA splicing of sarcomere genes. Circ Cardiovasc Genet 3, 138–146 (2010). 10.1161/CIRCGENETICS.109.904698

10 Liu, Y. et al. RNA-Seq identifies novel myocardial gene expression signatures of heart failure. Genomics 105, 83–89 (2015). 10.1016/j.ygeno.2014.12.002

11 Sweet, M. E. et al. Transcriptome analysis of human heart failure reveals dysregulated cell adhesion in dilated cardiomyopathy and activated immune pathways in ischemic heart failure. BMC Genomics 19, 812 (2018). 10.1186/s12864-018-5213-9

12 Heinig, M. et al. Natural genetic variation of the cardiac transcriptome in non-diseased donors and patients with dilated cardiomyopathy. Genome Biol 18, 170 (2017). 10.1186/s13059-017-1286-z

13 Zhao, S. & Zhang, B. A comprehensive evaluation of ensembl, RefSeq, and UCSC annotations in the context of RNA-seq read mapping and gene quantification. BMC Genomics 16, 97 (2015). 10.1186/s12864-015-1308-8

14 Steijger, T. et al. Assessment of transcript reconstruction methods for RNA-seq. Nat Methods 10, 1177–1184 (2013). 10.1038/nmeth.2714

15 David, J. K., Maden, S. K., Wood, M. A., Thompson, R. F. & Nellore, A. Retained introns in long RNA-seq reads are not reliably detected in sample-matched short reads. Genome Biol 23, 240 (2022). 10.1186/s13059-022-02789-6

16 Pardo-Palacios, F. J. et al. Systematic assessment of long-read RNA-seq methods for transcript identification and quantification. Nat Methods 21, 1349–1363 (2024). 10.1038/s41592-024-02298-3

17 Patowary, A. et al. Developmental isoform diversity in the human neocortex informs neuropsychiatric risk mechanisms. Science 384, eadh7688 (2024). 10.1126/science.adh7688

18 Glinos, D. A. et al. Transcriptome variation in human tissues revealed by long-read sequencing. Nature 608, 353–359 (2022). 10.1038/s41586-022-05035-y

19 Page, M. L. et al. A bioinformatic survey of RNA isoform diversity and expression across 9 GTEx tissues using long-read sequencing data. BMC Genomics 26, 1078 (2025). 10.1186/s12864-025-11919-w

20 Zhu, C. et al. Single-molecule, full-length transcript isoform sequencing reveals disease-associated RNA isoforms in cardiomyocytes. Nat Commun 12, 4203 (2021). 10.1038/s41467-021-24484-z

21 Pan, T. et al. Single-Cell Splicing Isoform Atlas of the Adult Human Heart and Heart Failure. Circulation 152, 1501–1514 (2025). 10.1161/CIRCULATIONAHA.125.074959

22 Haas, J. et al. Sarcomeric remodelling in human heart failure unraveled by single molecule long read sequencing. EMBO Mol Med 18, 824–845 (2026). 10.1038/s44321-025-00370-9

23 Frankish, A. et al. GENCODE: reference annotation for the human and mouse genomes in 2023. Nucleic Acids Res 51, D942–D949 (2023). 10.1093/nar/gkac1071

24 Pardo-Palacios, F. J. et al. SQANTI3: curation of long-read transcriptomes for accurate identification of known and novel isoforms. Nat Methods 21, 793–797 (2024). 10.1038/s41592-024-02229-2

25 Tardaguila, M. et al. SQANTI: extensive characterization of long-read transcript sequences for quality control in full-length transcriptome identification and quantification. Genome Res 28, 396–411 (2018). 10.1101/gr.222976.117

26 Amarasinghe, S. L. et al. Opportunities and challenges in long-read sequencing data analysis. Genome Biol 21, 30 (2020). 10.1186/s13059-020-1935-5

27 Workman, R. E. et al. Nanopore native RNA sequencing of a human poly(A) transcriptome. Nat Methods 16, 1297–1305 (2019). 10.1038/s41592-019-0617-2

28 Pan, Q., Shai, O., Lee, L. J., Frey, B. J. & Blencowe, B. J. Deep surveying of alternative splicing complexity in the human transcriptome by high-throughput sequencing. Nat Genet 40, 1413–1415 (2008). 10.1038/ng.259

29 Abugessaisa, I. et al. refTSS: A Reference Data Set for Human and Mouse Transcription Start Sites. J Mol Biol 431, 2407–2422 (2019). 10.1016/j.jmb.2019.04.045

30 Herrmann, C. J. et al. PolyASite 2.0: a consolidated atlas of polyadenylation sites from 3’ end sequencing. Nucleic Acids Res 48, D174–D179 (2020). 10.1093/nar/gkz918

31 Beaudoing, E., Freier, S., Wyatt, J. R., Claverie, J. M. & Gautheret, D. Patterns of variant polyadenylation signal usage in human genes. Genome Res 10, 1001–1010 (2000). 10.1101/gr.10.7.1001

32 Gruber, A. J. et al. A comprehensive analysis of 3’ end sequencing data sets reveals novel polyadenylation signals and the repressive role of heterogeneous ribonucleoprotein C on cleavage and polyadenylation. Genome Res 26, 1145–1159 (2016). 10.1101/gr.202432.115

33 Deviatiiarov, R. M. et al. An atlas of transcribed human cardiac promoters and enhancers reveals an important role of regulatory elements in heart failure. Nat Cardiovasc Res 2, 58–75 (2023). 10.1038/s44161-022-00182-x

34 Schoenauer, R. et al. EH-myomesin splice isoform is a novel marker for dilated cardiomyopathy. Basic Res Cardiol 106, 233–247 (2011). 10.1007/s00395-010-0131-2

35 Flam, E. et al. Integrated landscape of cardiac metabolism in end-stage human nonischemic dilated cardiomyopathy. Nat Cardiovasc Res 1, 817–829 (2022). 10.1038/s44161-022-00117-6

36 Subramanian, A. et al. Gene set enrichment analysis: a knowledge-based approach for interpreting genome-wide expression profiles. Proc Natl Acad Sci U S A 102, 15545–15550 (2005). 10.1073/pnas.0506580102

37 Liberzon, A. et al. Molecular signatures database (MSigDB) 3.0. Bioinformatics 27, 1739–1740 (2011). 10.1093/bioinformatics/btr260

38 Kohler, S. et al. The Human Phenotype Ontology in 2021. Nucleic Acids Res 49, D1207–D1217 (2021). 10.1093/nar/gkaa1043

39 Karlsson, M. et al. A single-cell type transcriptomics map of human tissues. Sci Adv 7 (2021). 10.1126/sciadv.abh2169

40 Nagy, E. & Maquat, L. E. A rule for termination-codon position within intron-containing genes: when nonsense affects RNA abundance. Trends Biochem Sci 23, 198–199 (1998). 10.1016/s0968-0004(98)01208-0

41 Patro, R., Duggal, G., Love, M. I., Irizarry, R. A. & Kingsford, C. Salmon provides fast and bias-aware quantification of transcript expression. Nat Methods 14, 417–419 (2017). 10.1038/nmeth.4197

42 Srivastava, A. et al. Alignment and mapping methodology influence transcript abundance estimation. Genome Biol 21, 239 (2020). 10.1186/s13059-020-02151-8

43 Mistry, J. et al. Pfam: The protein families database in 2021. Nucleic Acids Res 49, D412–D419 (2021). 10.1093/nar/gkaa913

44 Eddy, S. R. Accelerated Profile HMM Searches. PLoS Comput Biol 7, e1002195 (2011). 10.1371/journal.pcbi.1002195

45 Minegishi, S. et al. An Isoform of Nedd4-2 Plays a Pivotal Role in Electrophysiological Cardiac Abnormalities. Int J Mol Sci 18 (2017). 10.3390/ijms18061268

46 van Bemmelen, M. X. et al. Cardiac voltage-gated sodium channel Nav1.5 is regulated by Nedd4-2 mediated ubiquitination. Circ Res 95, 284–291 (2004). 10.1161/01.RES.0000136816.05109.89

47 Haywood, N. J. et al. Hypertrophic cardiomyopathy mutations in the calponin-homology domain of ACTN2 affect actin binding and cardiomyocyte Z-disc incorporation. Biochem J 473, 2485–2493 (2016). 10.1042/BCJ20160421

48 Wang, W. et al. Loss of Apelin exacerbates myocardial infarction adverse remodeling and ischemia-reperfusion injury: therapeutic potential of synthetic Apelin analogues. J Am Heart Assoc 2, e000249 (2013). 10.1161/JAHA.113.000249

49 Tatemoto, K. et al. Isolation and characterization of a novel endogenous peptide ligand for the human APJ receptor. Biochem Biophys Res Commun 251, 471–476 (1998). 10.1006/bbrc.1998.9489

50 Hiller, M., Huse, K., Platzer, M. & Backofen, R. Creation and disruption of protein features by alternative splicing -- a novel mechanism to modulate function. Genome Biol 6, R58 (2005). 10.1186/gb-2005-6-7-r58

51 Hiller, M., Huse, K., Platzer, M. & Backofen, R. Non-EST based prediction of exon skipping and intron retention events using Pfam information. Nucleic Acids Res 33, 5611–5621 (2005). 10.1093/nar/gki870

52 Love, M. I., Huber, W. & Anders, S. Moderated estimation of fold change and dispersion for RNA-seq data with DESeq2. Genome Biol 15, 550 (2014). 10.1186/s13059-014-0550-8

53 Vitting-Seerup, K. & Sandelin, A. IsoformSwitchAnalyzeR: analysis of changes in genome-wide patterns of alternative splicing and its functional consequences. Bioinformatics 35, 4469–4471 (2019). 10.1093/bioinformatics/btz247

54 Gilis, J., Vitting-Seerup, K., Van den Berge, K. & Clement, L. satuRn: Scalable analysis of differential transcript usage for bulk and single-cell RNA-sequencing applications. F1000Res 10, 374 (2021). 10.12688/f1000research.51749.2

