## Supplementary Figures for "A full-length isoform atlas of the non-failing and failing human heart"

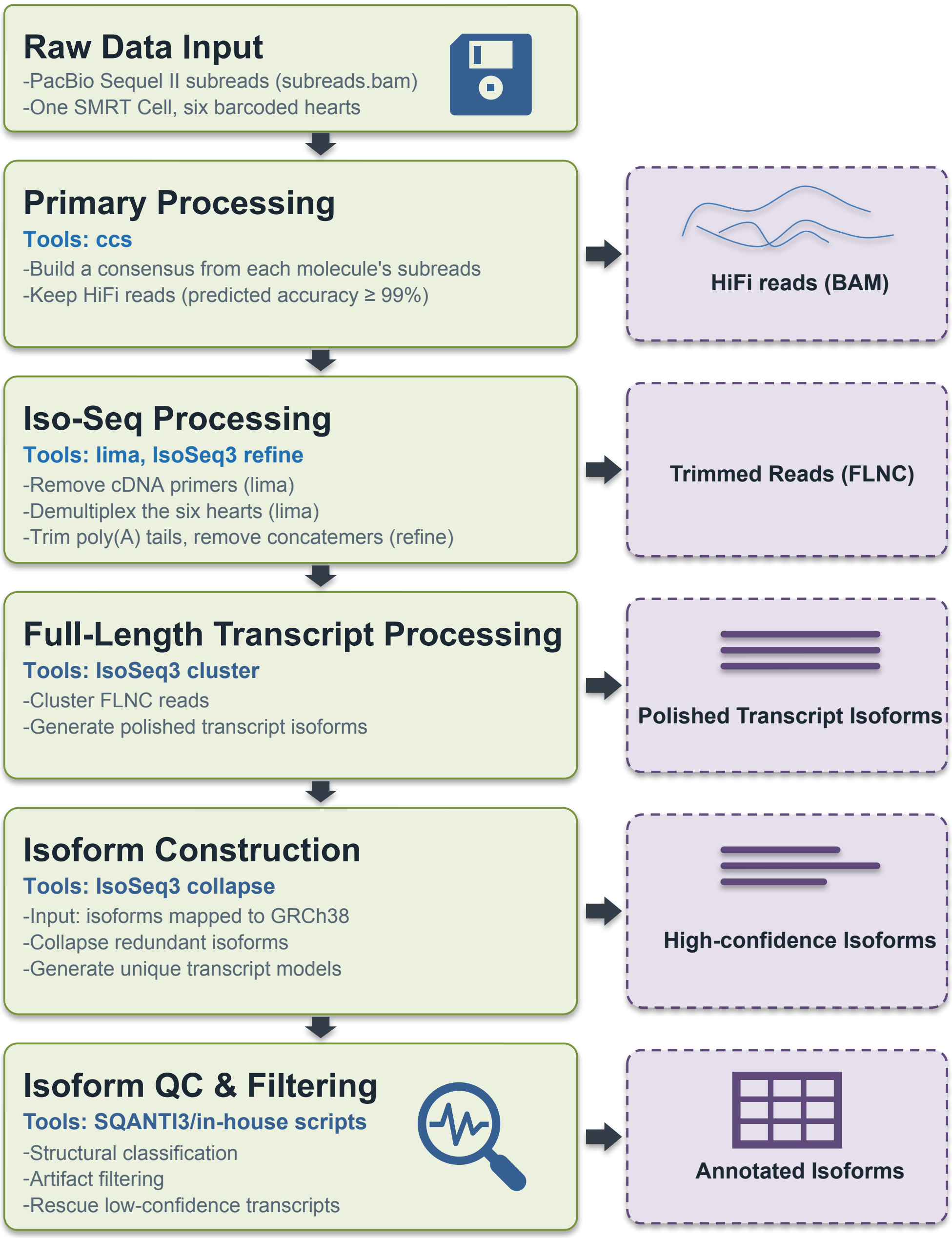


**Supplementary Figure 1. Long-read processing workflow.** PacBio Iso-Seq workflow for cardiac isoform catalog construction. Boxes on the left indicate processing stages, tools and operations; dashed boxes on the right show the corresponding output. Raw PacBio Sequel II subreads from one SMRT Cell carrying the six barcoded hearts undergo primary processing with ccs (v6.0.0), which builds a circular consensus of each molecule's subreads; the HiFi reads (predicted accuracy ≥ 99%) are kept. Lima removes the cDNA primers and demultiplexes the barcoded hearts, and IsoSeq3 refine trims poly(A) tails and removes concatemers, yielding full-length non-chimeric (FLNC) reads. IsoSeq3 cluster then groups the FLNC reads into polished transcript isoforms, and IsoSeq3 collapse merges redundant transcripts into nonredundant isoform models. Finally, SQANTI3 and in-house scripts perform structural classification against GENCODE v43, machine-learning-based artifact filtering, and rescue of low-confidence transcripts detected in at least 2 of 6 hearts, with at least 2 FLNC reads, 10 total reads, and support for at least one transcript end, yielding the curated cardiac isoform catalog.

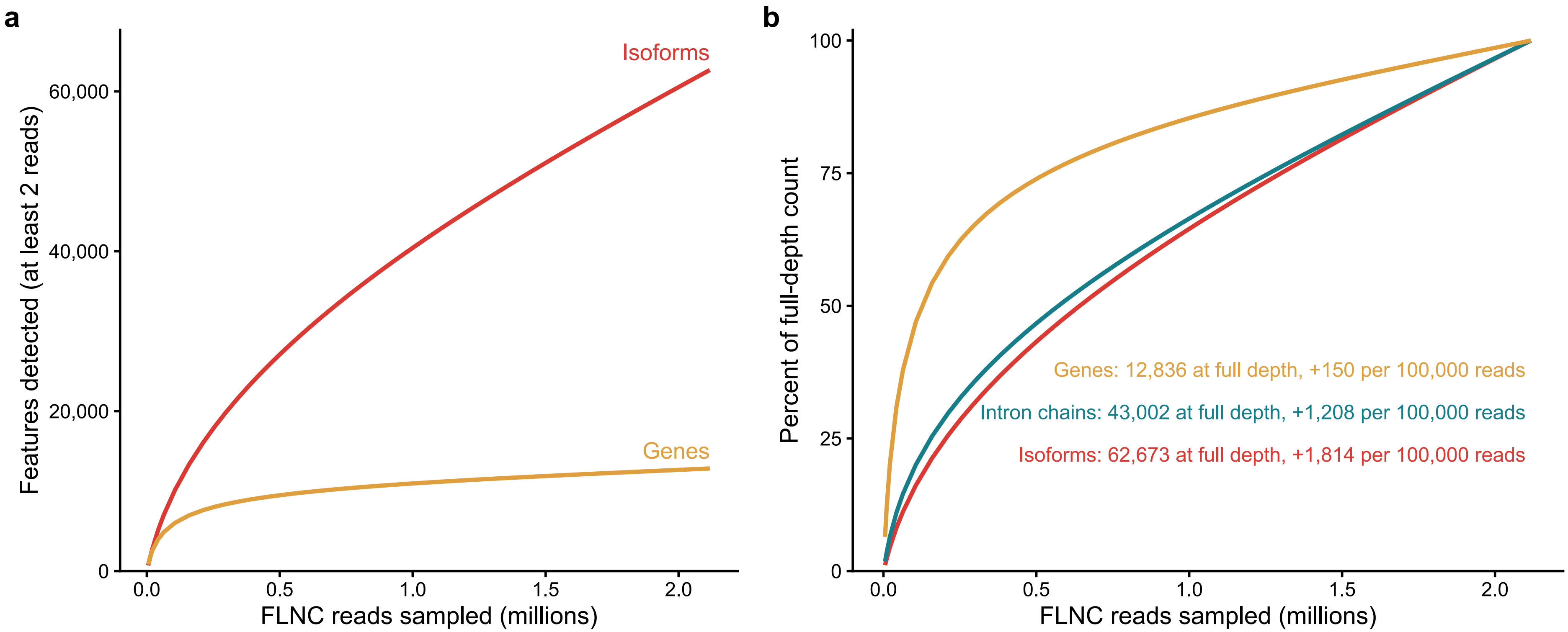


**Supplementary Figure 2. Gene discovery approaches saturation with sequencing depth, whereas isoform discovery does not.** Rarefaction of pooled FLNC reads from six hearts (n = 2,117,406), assigned to 62,673 collapsed isoforms. Curves show the analytically expected number of features supported by at least two sampled reads, matching the catalog inclusion criterion. Genes are SQANTI3 associated genes, with reads summed across their isoforms; intron chains are distinct ordered sets of splice junctions. a. Expected numbers of detected genes and isoforms as a function of sampled FLNC reads. b. Detection as a percentage of each feature's full-depth count, including intron chains. At 25% sequencing depth, 74% of genes were detected, compared with 47% of intron chains and 44% of isoforms. Even at full depth, discovery continued at approximately 150 genes, 1,200 intron chains, and 1,800 isoforms per additional 100,000 reads.

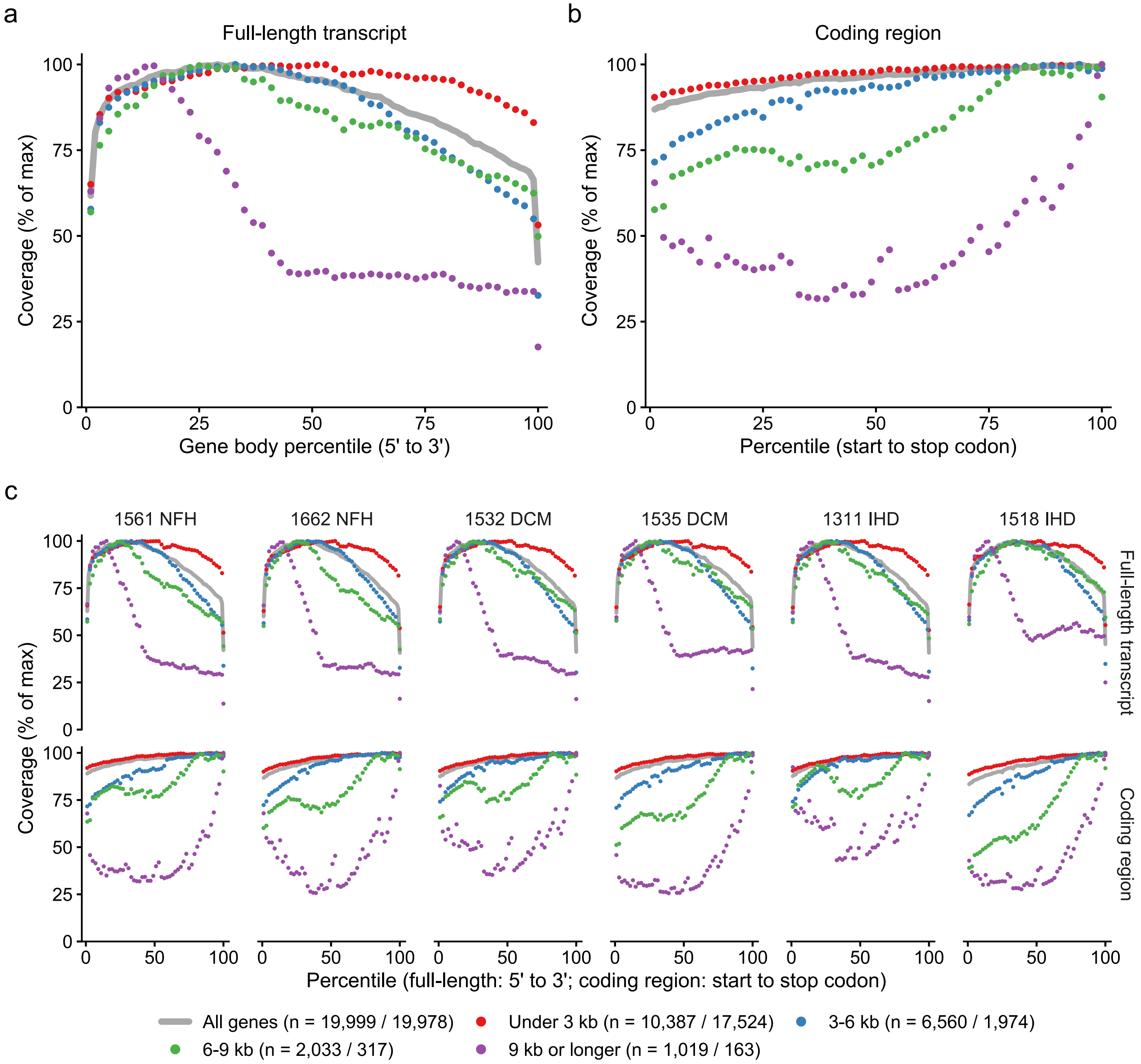


**Supplementary Figure 3. Read coverage along gene bodies, summed across genes and shown separately for each heart.** Coverage of FLNC reads along the GENCODE v43 canonical transcript of each protein-coding gene was sampled at 100 percentile positions, summed across genes within each length group, and shown as a percentage of the maximum. a. Coverage across the full-length transcript, from 5′ to 3′ (all genes, n = 19,999; under 3 kb, n = 10,387; 3-6 kb, n = 6,560; 6-9 kb, n = 2,033; 9 kb or longer, n = 1,019). b. Coverage across the coding region, from start to stop codon (all genes, n = 19,978; under 3 kb, n = 17,524; 3-6 kb, n = 1,974; 6-9 kb, n = 317; 9 kb or longer, n = 163). c. The corresponding profiles for each heart (NFH: 1561, 1662; DCM: 1532, 1535; IHD: 1311, 1518), showing similar coverage patterns across all six hearts. Because reads are summed across genes, these profiles are expression-weighted, with highly expressed genes contributing disproportionately; for example, TTN accounts for 53% of reads in the longest coding-length group. By contrast, Fig. 1d shows the median per-gene profile after normalizing each gene to its own peak, giving equal weight to every gene.

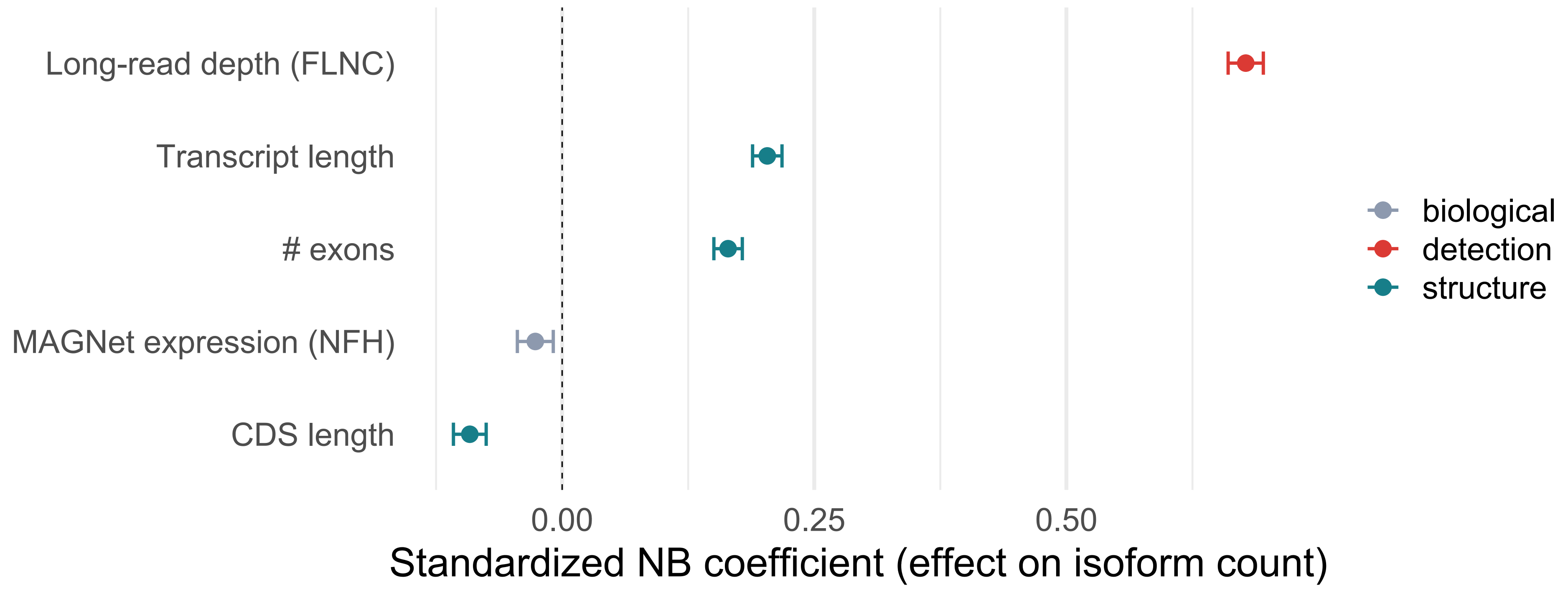


**Supplementary Figure 4. Long-read depth, rather than expression, is the main predictor of per-gene isoform counts.** Standardized coefficients from a multivariable negative-binomial model of curated long-read isoform counts per gene (n = 10,840 genes with nonzero FLNC depth and a MAGNet expression value). Predictors were log10-transformed and converted to z-scores: FLNC depth (reads summed across curated isoforms), canonical transcript length, exon count, mean MAGNet expression in non-failing hearts (CPM), and canonical coding sequence (CDS) length. Points show coefficients and bars show 95% confidence intervals; the dashed line marks zero. Colors distinguish detection (red), gene structure (teal) and biological expression (gray). FLNC depth was the dominant predictor ($\beta = 0.68$), whereas MAGNet expression had a near-zero coefficient ($\beta = -0.03$; partial Spearman $\rho = -0.05$ after controlling for depth). Fig. 1e shows the model after rarefaction to 10 reads per gene.

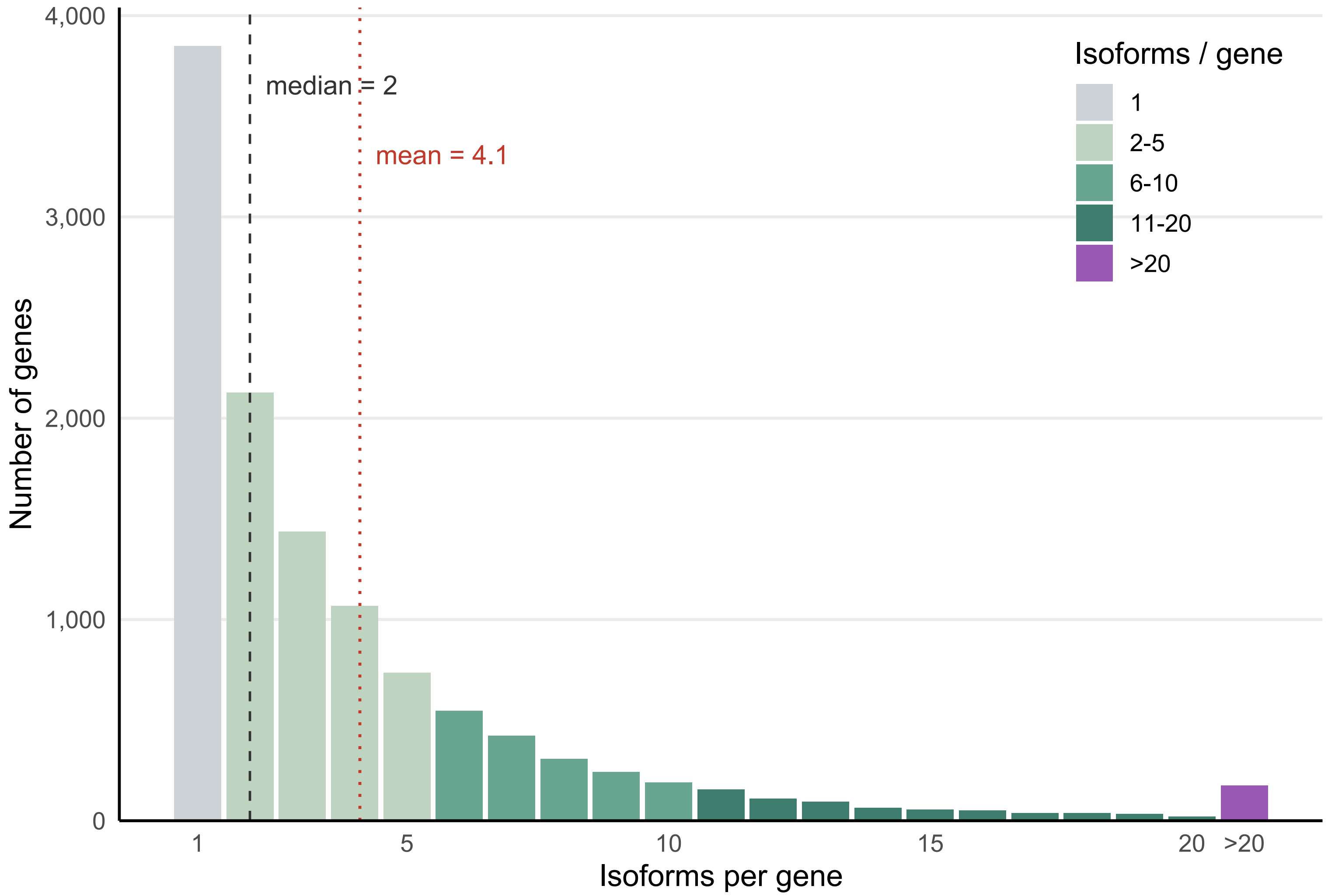


**Supplementary Figure 5. Number of curated isoforms per gene.** Histogram of curated isoform counts across the 11,760 genes in the catalog (48,232 isoforms). Counts from 1 to 20 are shown individually; genes with more than 20 isoforms are pooled. Bars are colored by diversity class: 1 isoform (3,848 genes, 32.7%), 2-5 (5,366, 45.6%), 6-10 (1,710, 14.5%), 11-20 (661, 5.6%) and more than 20 (175, 1.5%). The dashed gray and dotted red lines mark the median (2) and mean (4.1) isoforms per gene, respectively. Overall, 67% of genes (7,912) had multiple isoforms.

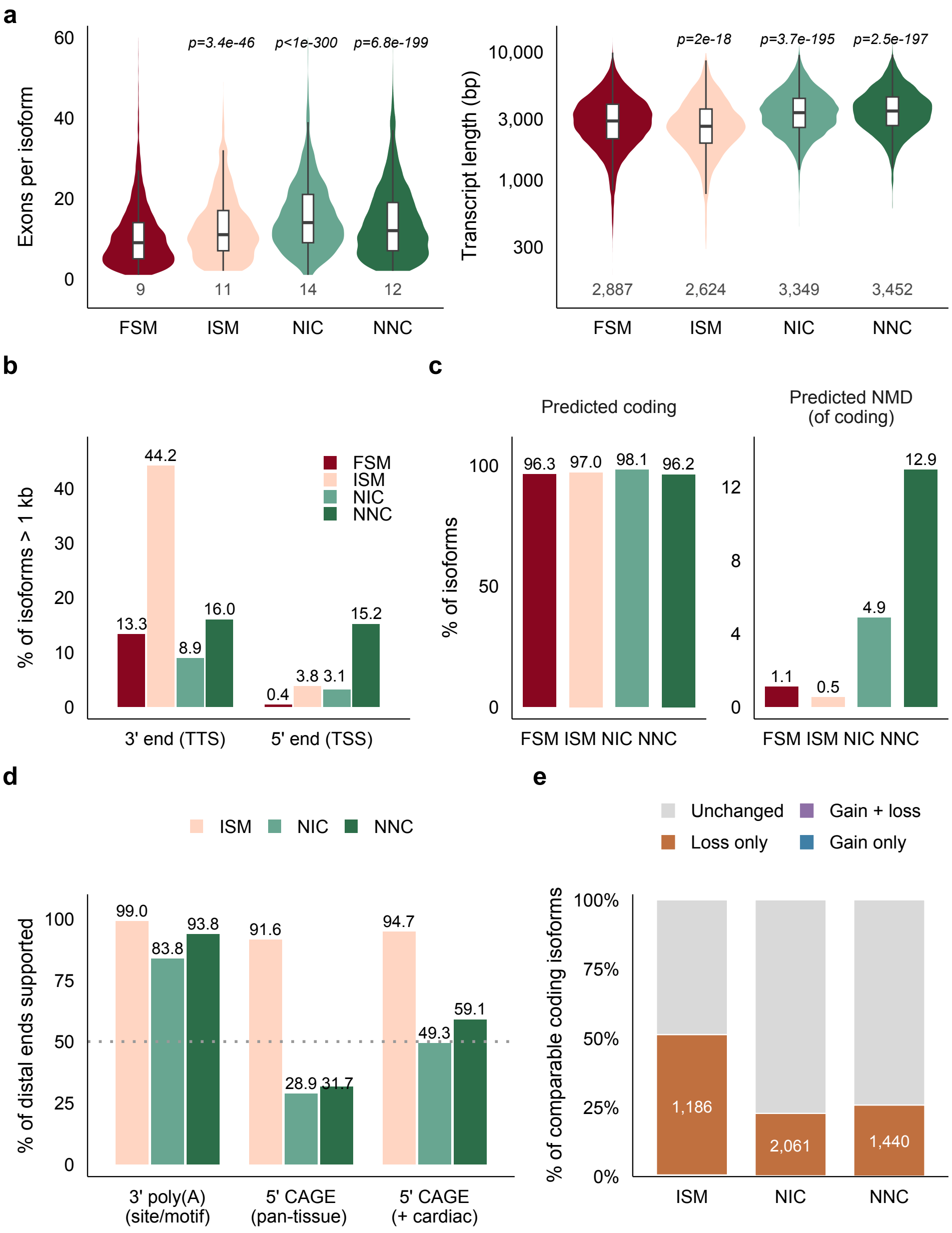


**Supplementary Figure 6. Structural features, coding potential, transcript-end support, and protein-domain content of novel and ISM isoforms relative to FSM.** a. Exon count (left) and transcript length (right, log10 scale) per isoform. Violins show the distribution, boxes indicate the median and interquartile range (IQR), with whiskers extending to 1.5 times the IQR. Numbers below violins indicate medians (exon count: FSM 9, ISM 11, NIC 14, NNC 12; length: 2,887, 2,624, 3,349, and 3,452 bp, respectively). P values are from two-sided Mann-Whitney tests against FSM. b. Percentage of isoforms with 3′ (TTS) or 5′ (TSS) ends more than 1 kb from the nearest annotated gene end, based on SQANTI3 distances. c. Percentage of isoforms predicted to encode proteins (left) and, among these, percentage predicted to be NMD-sensitive (right; note the different y-axis scales). d. Independent support for distal ends (> 1 kb from the annotated gene end) of ISM, NIC and NNC isoforms. Distal 3′ ends were assessed for overlap with a PolyASite 2.0 site or for a poly(A) signal motif (ISM, n = 1,102; NIC, n = 846; NNC, n = 966). Distal 5′ ends were assessed for overlap with a pan-tissue refTSS CAGE peak, with additional support from a human cardiac CAGE peak (GSE150736) within 50 bp (ISM, n = 95; NIC, n = 298; NNC, n = 916). The dotted line marks 50%. Panels b and d summarize the data in Fig. 4e,f. e. Pfam-domain content of comparable coding isoforms relative to the canonical GENCODE v43 protein for each gene (ISM, n = 2,338; NIC, n = 9,140; NNC, n = 5,607; isoforms whose protein and the canonical protein both lack Pfam domains were excluded). Isoforms are classed as unchanged, loss only, gain and loss, or gain only; gains were counted only when the gained domain sequence was absent from the canonical protein. Numbers in bars indicate loss-only isoform counts (ISM 1,186, NIC 2,061, NNC 1,440); gain-class counts are too small to label (gain and loss: 9, 11 and 6; gain only: 5, 15 and 4).

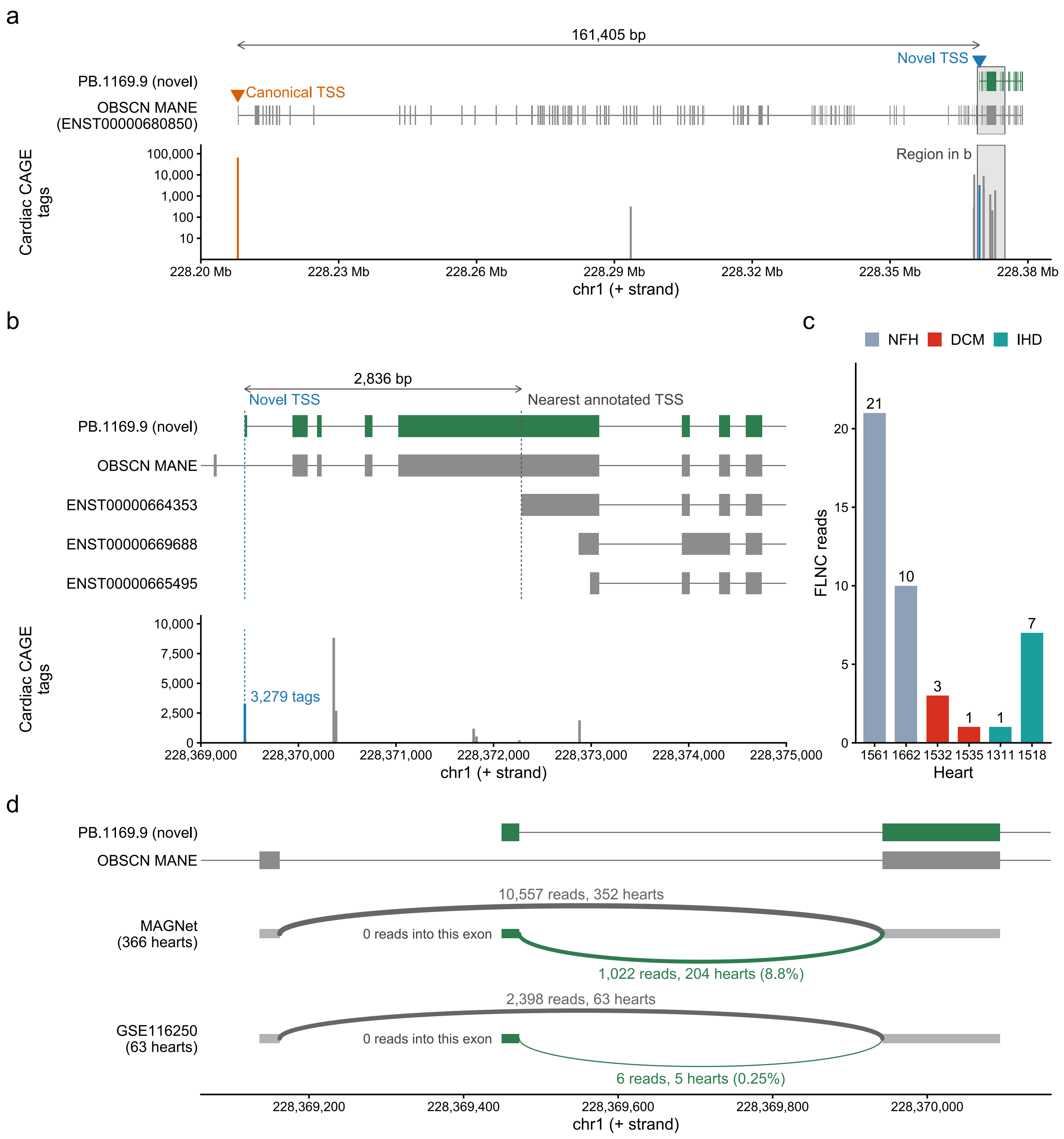


**Supplementary Figure 7. An unannotated internal promoter of OBSCN is supported by cardiac CAGE and short-read splice junctions.** a. The NNC isoform PB.1169.9 and the OBSCN MANE Select transcript (ENST00000680850) across the OBSCN locus, with sense-strand cardiac CAGE tag clusters (GSE150736; tags per cluster, log scale) shown below at each cluster's summit. PB.1169.9 starts 161,405 bp downstream of the canonical TSS (orange), within a CAGE cluster (blue). The shaded box marks the region shown in b. b. The 5′ end of PB.1169.9, the MANE transcript, and the three GENCODE v43 OBSCN transcripts with the nearest TSSs. Dashed lines mark the novel TSS and the nearest annotated TSS, 2,836 bp downstream. The 24-bp first exon of PB.1169.9 lies within a MANE intron and is spliced to the next MANE exon. Below, CAGE tags per cluster (linear scale); the cluster containing the novel TSS has 3,279 tags, with its summit 2 bp downstream of the long-read 5′ end. c. FLNC reads supporting PB.1169.9 in each heart (43 in total). d. Short-read splice junctions into the first exon shared by PB.1169.9 and the MANE transcript, in MAGNet (366 hearts) and GSE116250 (63 hearts). Shown are the summed unique junction reads across hearts and the number of hearts with at least one read for the junction from the novel first exon (green; its share of the reads from the two junctions shown in parentheses) and the junction from the preceding MANE exon (gray). Arc width scales with log10 read count; junctions into the same exon with 9 or fewer reads are not shown. No junction in either cohort ends at the novel first exon.

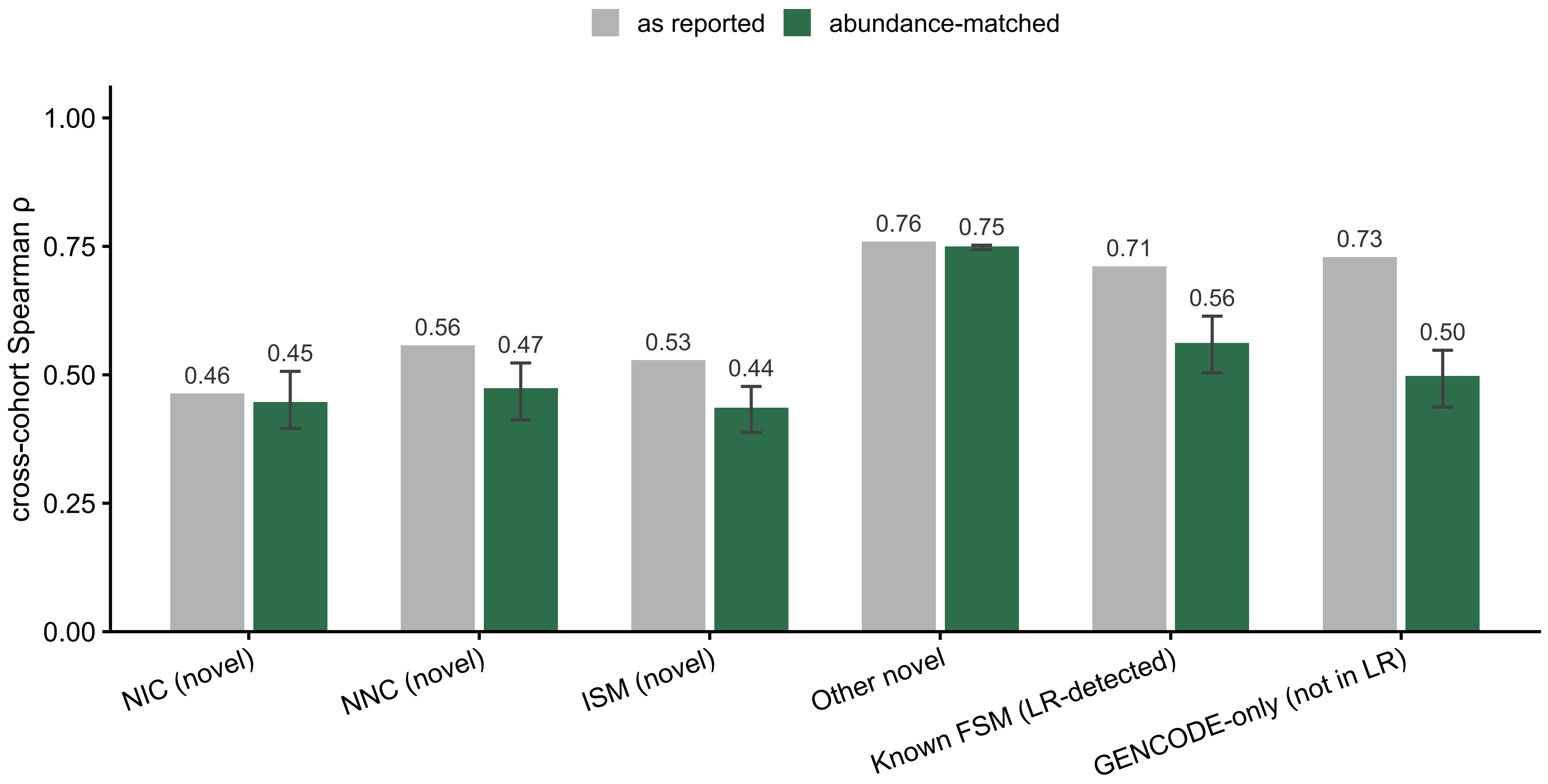


**Supplementary Figure 8. Cross-cohort reproducibility of isoform expression after abundance matching.** Mean transcript TPM was calculated separately in MAGNet (n = 366) and GSE116250 (n = 63), and Spearman correlations (ρ) between cohort means were computed for transcripts expressed in both cohorts: NIC (n = 7,396), NNC (n = 5,373), ISM (n = 1,407), other novel (fusion, genic, antisense, and intergenic; n = 576), known FSM (GENCODE transcripts matched to curated long-read isoforms; n = 16,256), and GENCODE-only (not detected in long-read data; n = 177,565). Gray bars show correlations across all transcripts in each category (Fig. 5f). Green bars show correlations after abundance matching: each category was randomly down-sampled to the same abundance distribution (0.25-log10 bins of the geometric mean of the two cohort means), retaining the minimum count across categories in each bin (571 transcripts per category). Green bars represent the mean of 300 random down-samplings and error bars indicate the 2.5th to 97.5th percentiles. The interval for other novel isoforms is narrow because nearly all 576 transcripts are included in each draw. After matching, correlations for NIC (0.45), NNC (0.47), and ISM (0.44) were comparable to the correlation for GENCODE-only transcripts (0.50), with overlapping intervals.

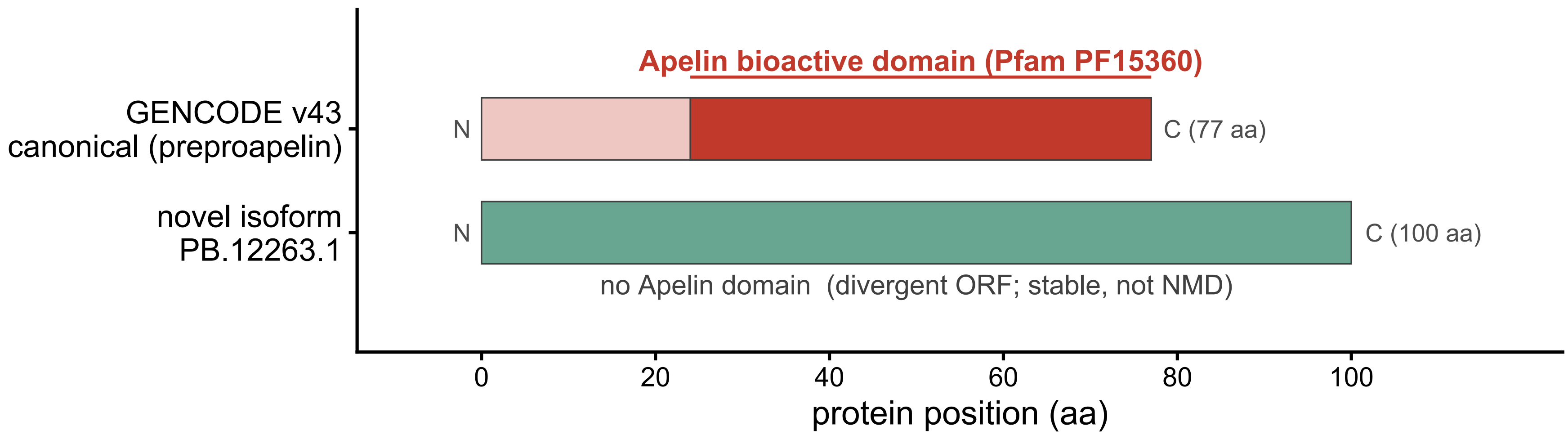


**Supplementary Figure 9. A divergent APLN reading frame lacks the bioactive Apelin domain.** Protein schematics of canonical GENCODE v43 APLN preproapelin (ENST00000429967; 77 aa) and the predicted ORF of NNC long-read isoform PB.12263.1 (100 aa). The canonical Apelin domain (Pfam PF15360; residues 25-77, dark red), from which bioactive apelin peptides are cleaved, is absent from PB.12263.1 because its reading frame diverges from that of preproapelin (frameshift-divergent loss). PB.12263.1 was not predicted to be NMD-sensitive.

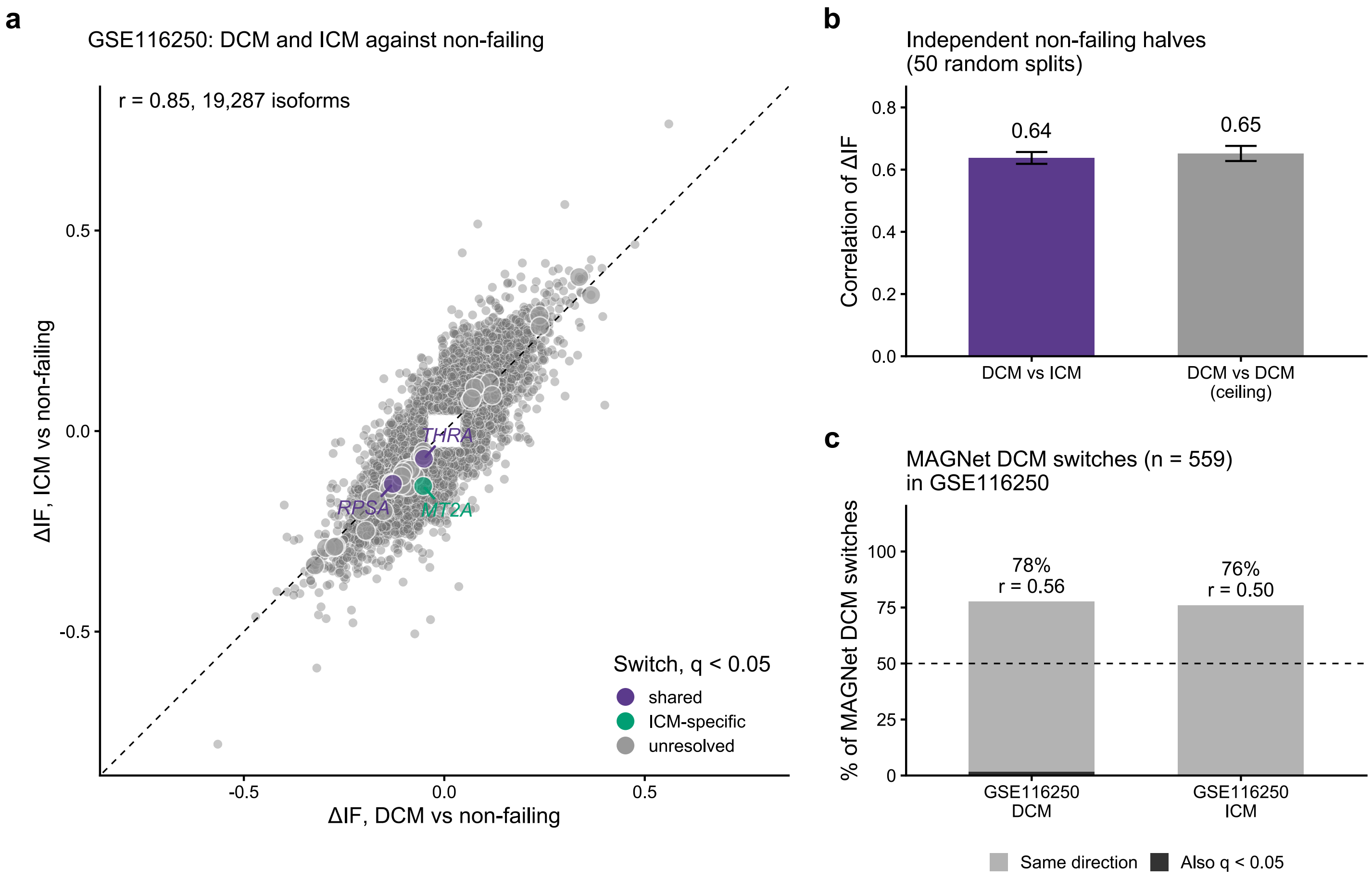


**Supplementary Figure 10. Isoform changes in dilated and ischemic cardiomyopathy are largely shared.** Isoform switching in GSE116250 was analyzed using a single IsoformSwitchAnalyzeR model (satuRn test) across 14 non-failing, 36 DCM, and 13 ischemic cardiomyopathy (ICM) hearts. Transcript abundance was quantified with Salmon against GENCODE v43 augmented with 18,637 long-read isoforms. Age, sex, and surrogate variables were included as covariates, with three contrasts: DCM vs non-failing, ICM vs non-failing, and ICM vs DCM. a. Change in isoform fraction (ΔIF) in DCM versus non-failing hearts plotted against ΔIF in ICM versus non-failing hearts for 19,287 isoforms with |ΔIF| ≥ 0.05 in either disease. Disease effects were strongly correlated (Pearson r = 0.85). Larger points mark switches significant in at least one contrast (q < 0.05 and |ΔIF| ≥ 0.05; 35 in DCM and 4 in ICM, 37 unique isoforms). Two were significant in both diseases in the same direction (RPSA and THRA); one was ICM-specific and significant in the direct ICM vs DCM contrast (MT2A). The other 34 were unresolved, significant in one disease but not in the direct comparison. The direct ICM vs DCM contrast identified 10 significant isoforms across 8 genes. b. Correlation of ΔIF between DCM and ICM using separate halves of the non-failing hearts as references (7 each; purple, r = 0.64), compared with the correlation between two independent groups of 13 DCM hearts (gray, r = 0.65), representing the expected agreement if the disease effects were identical. Bars show means across 50 random splits; error bars indicate SD. Isoforms are as in panel a. c. Directional concordance of 559 MAGNet DCM isoform switches testable in GSE116250 (of 652 total; q < 0.05, |ΔIF| ≥ 0.05): 78% in GSE116250 DCM and 76% in ICM. r indicates the correlation of ΔIF between cohorts. Dark segments indicate switches also significant in GSE116250 (9 in DCM; 1 in ICM). Dashed line marks 50% expected by chance.

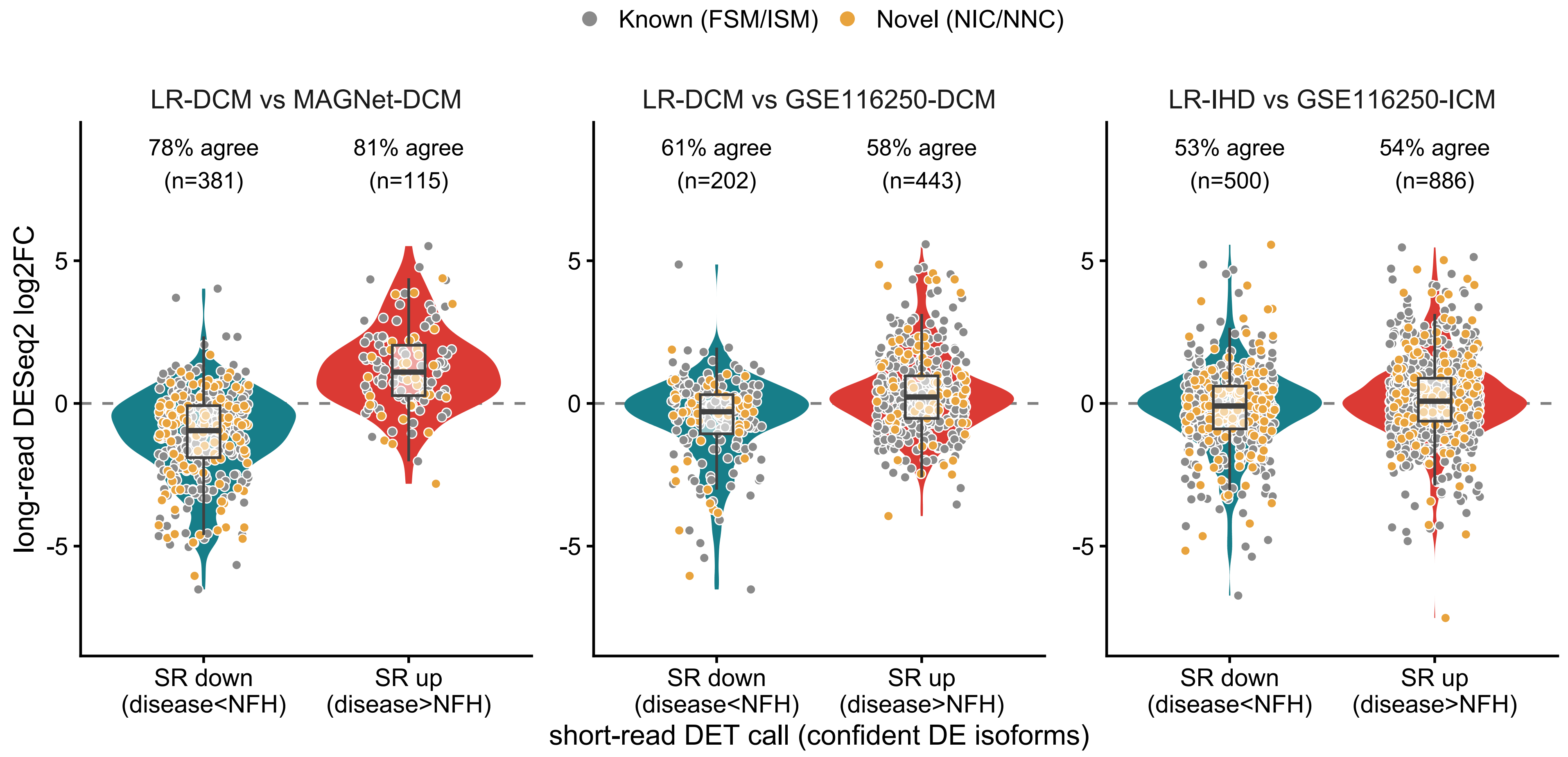


**Supplementary Figure 11. Directional concordance of long-read differential expression with short-read cohorts.** Because the long-read cohort included only two hearts per phenotype, reproducibility was assessed by direction rather than fold-change magnitude. For each isoform confidently differential in a short-read cohort (adjusted $p < 0.05$ and $|\log2FC| > 1$), the long-read DESeq2 log2FC (disease vs non-failing) is plotted by short-read (SR) direction: SR down or SR up. The dashed line marks zero; concordant isoforms fall below zero for SR down and above zero for SR up. Violins show distributions, boxes show medians and interquartile ranges, and points represent individual isoforms (gray, known FSM/ISM; gold, novel NIC/NNC). Labels at the top of each panel give the percentage of isoforms with matching directions and the number tested (n). a. Long-read DCM vs MAGNet DCM. b. Long-read DCM vs GSE116250 DCM. c. Long-read IHD vs GSE116250 ischemic cardiomyopathy (ICM). Agreement was highest against MAGNet (78.6% of 496 isoforms; 78.9% of 147 novel isoforms) and lower against GSE116250 (DCM, 58.6% of 645; ICM, 53.6% of 1,386), but exceeded chance in both.

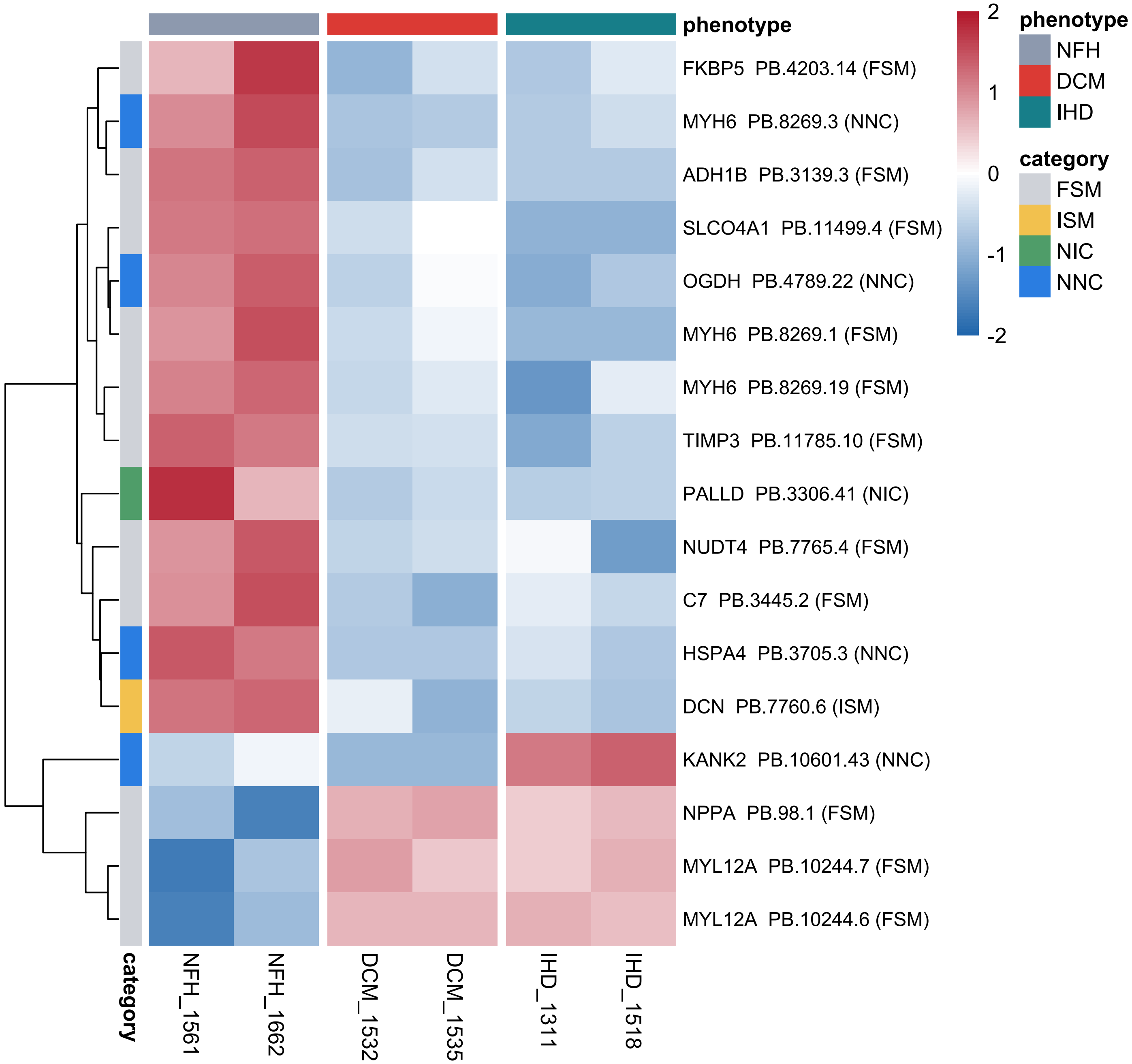


**Supplementary Figure 12. Isoforms differentially abundant across phenotypes in the long-read cohort.** Heatmap of the 17 isoforms with significant omnibus differential abundance across the three phenotypes (DESeq2 likelihood-ratio test comparing the full phenotype model with an intercept-only model; Benjamini-Hochberg adjusted $p < 0.05$). Testing included 17,945 curated isoforms with at least 10 total FLNC reads and reads in at least 2 of the 6 hearts. Columns are the six hearts grouped by phenotype (NFH: 1561, 1662; DCM: 1532, 1535; IHD: 1311, 1518). Rows are labeled by gene, isoform and structural category (left color bar: 11 FSM, 1 ISM, 1 NIC, 4 NNC) and ordered by hierarchical clustering. Thirteen isoforms were higher in both NFH hearts than in disease hearts; KANK2 PB.10601.43 was highest in IHD; and NPPA and two MYL12A isoforms were higher in DCM and IHD.
